# Dissection of ergosterol metabolism reveals a pathway optimized for membrane phase separation

**DOI:** 10.1101/2025.01.05.631358

**Authors:** Israel Juarez-Contreras, Laura J.S. Lopes, Jamie Holt, Lorena Yu-Liao, Katherine O’Shea, Jose Ruiz-Ruiz, Alexander Sodt, Itay Budin

**Affiliations:** Department of Chemistry and Biochemistry, University of California San Diego, La Jolla, CA 92093; Unit on Membrane Chemical Physics, Eunice Kennedy Shriver National Institute of Child Health and Human Development; 29 Lincoln Drive, Bethesda, MD 20892

## Abstract

Sterols are among the most abundant lipids in eukaryotic cells, yet are synthesized through notoriously long metabolic pathways. It has been proposed that the molecular evolution of such pathways must have required each step to increase the capacity of its product to condense and order phospholipids. Here we carry out a systematic analysis of the ergosterol pathway that leverages the yeast vacuole’s capacity to phase-separate as a predictive biophysical readout for each intermediate. In the post-synthetic steps specific to ergosterol biosynthesis, we find that successive modifications act to oscillate ordering capacity, settling on a level that supports phase separation while retaining fluidity of the resulting domains. Simulations carried out with each intermediate showed how conformers in the sterol’s alkyl tail are capable of modulating long-range ordering of phospholipids, which could underlie changes in phase behavior. Our results indicate that the complexity of sterol metabolism could have resulted from the need to balance lipid interactions required for membrane organization.

## Introduction

Sterols are polycyclic triterpenoid lipids that are hallmarks of eukaryotic cell membranes. Canonical structural roles for sterols are based on their ability to condense the hydrocarbon chains of neighboring phospholipids, thereby ordering bilayers in a concentration-dependent manner (*1*). While this function might be universal, it does not explain the chemical diversity in sterol metabolism among eukaryotic lineages (*2*). Synthesis of sterols is metabolically complex, with the most common pathways featuring 10 or 11 distinct steps, several of which are oxygen dependent. The recent discovery of protosterols in the fossil record corresponding to each of the crown group eukaryotes supports a model in which the diversification of sterol metabolism that occurred during the early evolution of eukaryotes, likely alongside increasing oxygen levels during the Tonian period (*3*). While the initial steps for each pathway are similar, their subsequent diversification suggests disparate selective pressures during eukaryotic evolution (*4*). In the case of ergosterol, the most common fungal sterol, early steps are shared with cholesterol biosynthesis in metazoans, while later steps are unique and differentiate the pathway’s final product (Fig. 1A).

**Fig. 1.**
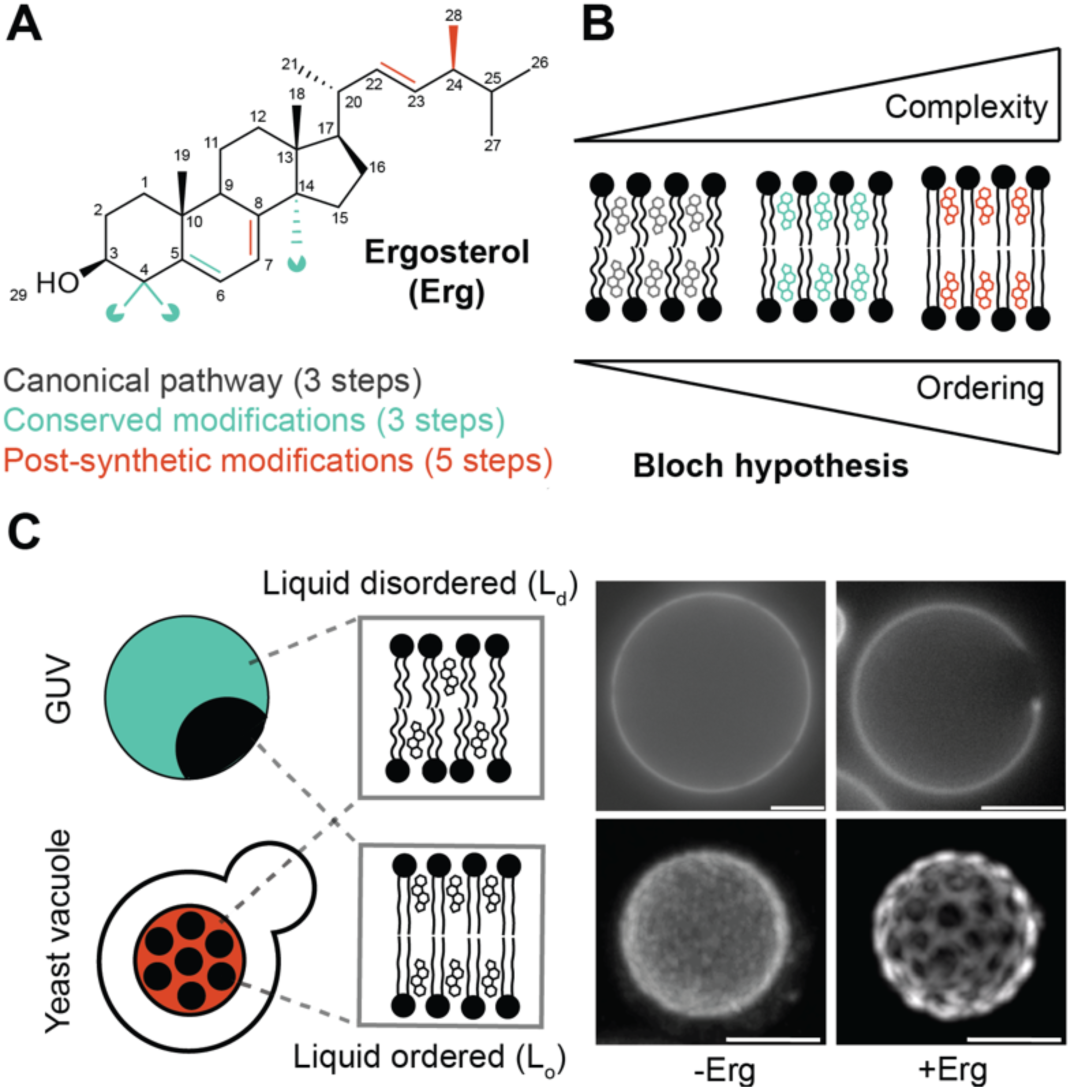
Biosynthesis of sterols and their role in membrane phase separation. (**A**), Chemical structure of the fungal sterol ergosterol, with highlighted modifications grouped according to their occurrence during its metabolism. The canonical sterol pathway yields the ring system, conserved modifications among fungi and animals result in demethylation of α-face carbons, while post-synthetic modifications exclusive to ergosterol primarily remodel the alkyl chain. (**B**), Schematic of the Bloch hypothesis, which proposes that the capacity for sterols to condense and order phospholipids rose continuously along the pathway. (**C**), Liquid phase separation between L_d_ and L_o_ domains is a sterol-dependent membrane property that can be observed in both synthetic systems (giant unilamellar vesicle, GUV) and in cells (the yeast vacuole). In both GUVs and vacuoles, changes in ergosterol concentrations drive the loss or appearance of L_o_/L_d_ phase separation. Shown are 3:1 DOPC:DPPC GUVs prepared with 0 (left) and 20 mol % (right) ergosterol (top) and yeast vacuoles from an ergosterol depleted strain (*ERG9* knockdown, left) compared to WT (right). Scale bars, 5 µm.

The complexity of sterol biosynthesis has long motivated models for understanding the evolution of metabolic pathways. Based on vesicle permeability, NMR, and microviscosity measurements in mixtures with phosphatidylcholine (PC), Konrad Bloch and colleagues first observed that cholesterol has a stronger ordering capacity than its first precursor lanosterol, with demethylated cholesterol precursors in the pathway showing intermediate effects (*5–7*). These experiments led to the proposal that sterol metabolism has evolved primarily to increase membrane ordering for improved barrier functions of cell membranes (*8*). In the Bloch hypothesis, each pathway intermediate is expected to show superior ordering capability, thereby explaining their sequential adoption during the evolution of the pathway. Such an optimization is best understood through changes in the bifacial topology of the sterol ring system after lanosterol synthesis: one plane (the α-face) is demethylated three times for optimal van der Waals interactions with neighboring phospholipid chains, while the other (the β-face) retains two vestigial methyl groups that contribute to its orientation in the bilayer (*9, 10*). These demethylations are conserved across pathways. However, subsequent steps that are unique to each pathway have not been explained by the Bloch hypothesis, such as the generation of the methylated tail in ergosterol (*11*) and plant phytosterols (*12*). These sterols have been observed to show inferior ordering capabilities compared to cholesterol (*8, 13*), leading to models in which factors, such as temperature (*8*), desiccation (*14*), ultraviolet irradiation (*3*), could have instead driven their evolution.

The Bloch hypothesis was formulated before the discovery that sterols are required to support lateral membrane heterogeneity, a process that depends on their interaction with multiple lipids. When included in mixtures of high and low melting temperature phospholipids or sphingolipids, sterols can drive membrane phase separation into liquid ordered (L_o_) and liquid disordered (L_d_) domains (Fig. 1B) (*15*). Colloquially termed lipid rafts, this emergent property has been well characterized in model membrane systems, such as giant unilamellar vesicles (GUVs) (*16*). Without sterols, high melting temperature lipids can still demix, but form gel-like, solid domains whose reduced dynamics might preclude biological functions (*17*). While sterols are necessary for fluid membrane phase separation, the specific structural requirements for this process have not been elucidated. The α-face methylated sterol lanosterol does not support domains in model membranes (*18*), indicating that ordering capacity is required for domain formation. However, even late-stage intermediates, such as 7-dehydrocholesterol that accumulate in patients with Smith-Lemli-Opitz syndrome, can also show altered domain forming capacity (*19*). A set of sterol derivatives, classified as inhibitor sterols, has also been observed to prevent liquid domain formation, instead disrupting phase separation (*20*) or favoring the formation of solid domains (*18*). Together, these studies suggest that membrane phase separation is sensitive to specific chemical features of sterols. More systematic analyses of sterol metabolites have been hampered by their limited commercial and synthetic availability.

Though best understood in model systems, ordered membrane domains are also thought to be relevant for molecular organization *in vivo*. In mammalian cells, domains are small and dynamic, making systematic characterization challenging (*21*). An alternative model system is the vacuole of yeast cells (Fig. 1C). Upon nutritional stress or transition of cells into the stationary growth stage, vacuole membranes organize into micron-scale domains (*22*). These structures can act as internalization sites for neutral lipid stores that help cells survive under these conditions (*23*). Vacuole domains show hallmarks of phase separation in model membranes: fluid-like mobility, a tendency to coalesce upon collision, and a characteristic miscibility temperature above which they reversibly dissolve (*24*). Domain formation during early stationary stage growth corresponds to an increase in membrane ergosterol composition, alongside that of high melting temperature sphingolipids (*25*), and domains from isolated vacuoles are sensitive to sterol depletion (*26*), as they are in vesicles (*27*). Genetic and chemical depletion of ergosterol from cells also prevents vacuole domains from forming in cells (*22*). Alterations in the abundance structure of vacuole sphingolipids modulate domain structure (*25*), suggesting that a balance of distinct lipid species is required for robust phase separation of liquid domains.

Here, we take advantage of yeast vacuoles to investigate the sterol requirements for membrane phase separation. Motivated by the Bloch hypothesis, we asked whether successive intermediates in the ergosterol pathway exhibit improved capacities to form ordered domains. Extraction of sterol intermediates from strains allowed us to identify these features and measure their interactions with phospholipids, which was further explored with all-atom simulations. Together, these data uncover a pathway that does not maximize the condensation capacity, but rather balances it to support phase separation of fluid domains.

## Results

### Genetic modulation of early-stage modifications and effects on phase separation

To directly test the function of intermediates, we generated yeast strains to systematically modify sterol composition in cells. Early stages of the pathway, shared with cholesterol synthesis in metazoans, form the initial cyclized product lanosterol and its subsequent demethylation into zymosterol. These steps include the condensation of two farnesyl pyrophosphate into squalene by Erg9, squalene oxidation into 2,3-oxidosqualene by Erg1, and the formation of lanosterol by the cyclase Erg7. Lanosterol is demethylated by Erg11 into 4,4-dimethylcholesta-8,14,24-trienol (or follicular fluid meiosis-activating sterol, FF-MAS), reduced into 4,4-dimethylzymosterol by Erg24, and again demethylated by the coordinated activities of the Erg25, Erg26, and Erg27 into zymosterol. Because these steps are all essential for yeast viability, we used a promoter replacement strategy to place expression of their coding genes under control of supplemental doxycycline (P_tetoff_) or methionine (P_MET_) (Fig. 2A). The resulting strains are summarized in Table S1.

**Fig. 2.**
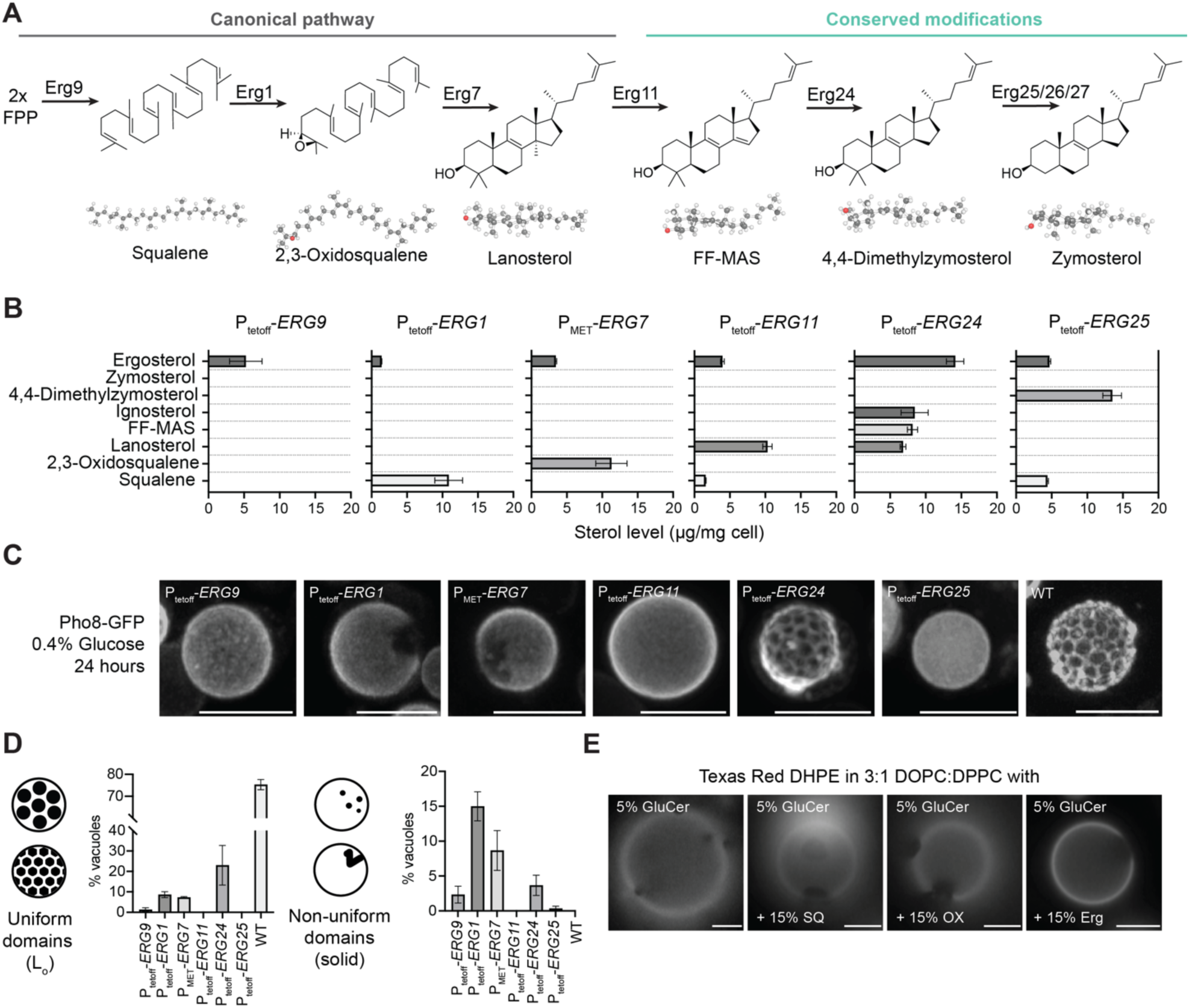
Early-stage intermediates show limited capacity for L_o_/L_d_ phase separation. (**A**), Schematic of canonical sterol-forming steps and conserved modifications, shared between ergosterol and cholesterol synthesis. (**B**), Sterol composition of knockdowns in each early-stage step as measured by gas chromatography-mass spectrometry (GC-MS). N = 3 independent cultures; error bars = SD. (**C**), Representative micrographs for each strain. Among mutants, only ERG24 knockdown still yields robust L_o_ domain formation, likely because of increased residual ergosterol content. Scale bars, 5 µm. Wider field images with multiple cells are shown in Fig. S1A. (**D**), Quantification of uniform (L_o_) and non-uniform (solid) domain formation frequency in stationary phase cells. Gel-like domains are abundant in mutants that accumulated squalene or 2,3-oxidosqualene. N = 3 independent cultures with n > 100 cells each; error bars = SEM. **E.** Early stage uncyclized intermediates squalene (SQ) and 2,3-oxidosqualene (OX) at 15 mol % cause expansion of solid or gel-like domains formed by 5 mol % of the glycosphingolipid glucosylceramide (GlcCer) in 3:1 DOPC:DPPC GUVs. In contrast, ergosterol supports fluid L_o_/L_d_ domains (right). The presence of multiple domains in GUVs allowed to equilibrate below the mixing temperatures indicates non-fluid domains that do not coalesce. Scale bars, 5µm.

We hypothesized that repression of specific *ERG* genes would allow sufficient expression for viability, while strongly reducing the accumulation of the subsequent biosynthetic product. To test this, we measured sterol compositions of whole cells grown under vacuole domain-forming conditions under continual repression. The resulting profiles (Fig. 2B) changed from wild-type (WT) cells as expected based on the genes targeted, with an accumulation of the upstream intermediate in the pathway. For *ERG24* knockdown (P_tetoff_-*ERG24*), accumulation of the expected intermediate FF-MAS was accompanied by a derivative (ignosterol) previously observed in cells treated with antifungal Amorolfine (*28*). All strains retained residual ergosterol, presumably supporting their viability, but at levels <10% of WT cells except for P_tetoff_-*ERG24*.

We next tested the effects of accumulating early-stage intermediates on ordered domain formation in the yeast vacuole (Fig. 2C, S1A). Growth under metabolic restriction (0.4% glucose) drives vacuole membrane phase separation in most cells imaged using the L_d_ marker Pho8-GFP. All early-stage knockdowns showed reductions in domain formation, supporting the essential role of ergosterol in this process (Fig. 2D). Compared to *ERG9* knockdown cells, which contained very few phase-separated vacuoles, *ERG1* and *ERG7* knockdowns that accumulate linear precursors (squalene and 2,3-oxidosqualene) showed enhanced domain formation. However, these were predominantly irregularly shaped and resisted coarsening upon collision, suggesting they represented solid or gel-like domains previously observed in yeast sphingolipid mutants (*25*). Solid domains were observed in half and two-thirds of all phase-separated vacuoles in *ERG7* and *ERG1* knockdown cells, respectively. The only early-stage knockdown to support robust uniform domain formation was that for *ERG24*, which accumulated FF-MAS and its derivatives. However, *ERG24* knockdown cells also contained higher amounts of ergosterol than other strains, suggesting that its phenotype was not based on the accumulated intermediates themselves. Consistent with this hypothesis, we found that FF-MAS itself does not support lateral phase separation in GUVs (Fig. S2).

Given that *ERG1* and *ERG7* knockdown cells showed levels of domain formation not observed in subsequent pathway steps, we asked if linear sterol precursors affect domain membrane formation in GUVs. In ternary mixtures with 60% di-oleoyl PC (DOPC) and 20% di-palmitoyl PC (DPPC), we observed that both squalene and 2,3-oxidosqualene inhibit formation of Lo domains in mixtures with ergosterol (Fig. S3). However, in vacuoles, the accumulation of these intermediates caused formation of irregular gel-like domains, which are associated with high melting temperature glycosphingolipids (*29*). We therefore tested if linear sterol precursors promoted phase separation in membranes containing the glucosyl-ceramide (GlcCer), which resembles the predominant vacuole yeast sphingolipid inositol phosphoceramide (IPC). We observed that 15% of squalene or 2,3-oxidosqualene cause gel-like domains to enlarge in GUVs containing 5% GlcCer (Fig. 2E). In contrast, ergosterol solubilized GlcCer into fluid L_o_ domains, analogous to the state of WT vacuoles. Thus, early sterol precursors could have roles in promoting membrane heterogeneity, but overall lack the capacity to support liquid (L_o_/L_d_) phase separation as ergosterol does.

### How domain formation capacity arises along the post-synthetic steps

We next explored precursors that arise in the late-stage pathway, termed post-synthetic steps by Bloch (*8*), which are specific to ergosterol metabolism (Fig. 3A). In this part of the pathway, the alkyl tail is methylated (Erg6) and its double bond moved to the C22(23) position through the successive actions of a desaturase (Erg5) and a reductase (Erg4). On the B-ring, the isomerase Erg2 converts the Δ^8^ double bond to a Δ^7^ one, and the desaturase Erg3 introduces an additional double bond at Δ^5^. These actions lead to a final product whose B-ring is diunsaturated (Δ^5,7^) and a 6-carbon tail that is monounsaturated at C22 and methylated at C24(28).

**Fig. 3.**
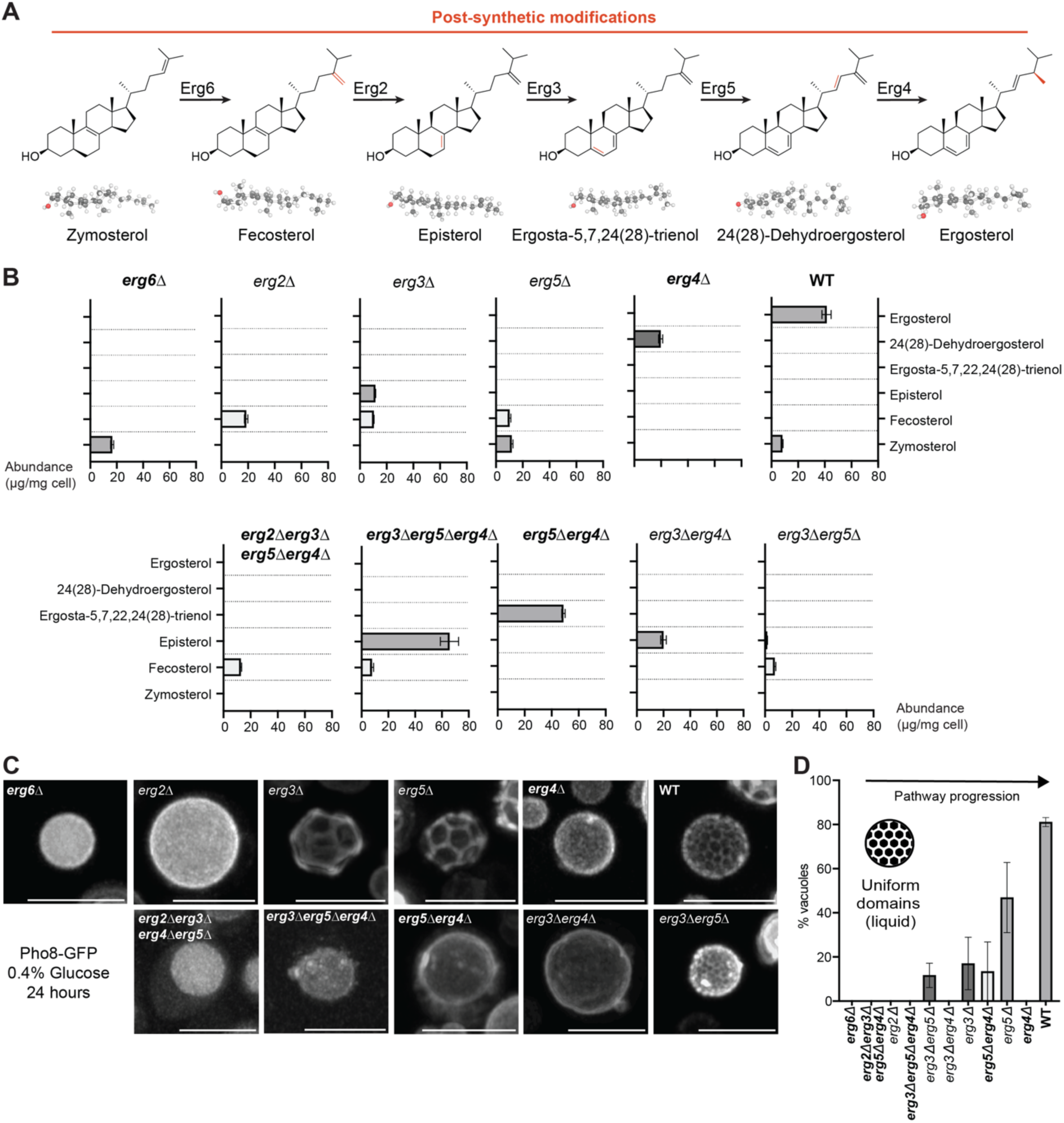
Post-synthetic sterol intermediates show a non-monotonic increase in ability to support membrane phase separation. (**A**), Schematic of post-synthetic steps transforming zymosterol into ergosterol. (**B**), Sterol composition of mutants that isolate each of the post-synthetic steps as measured. Mutants in bold predominantly produce an ergosterol intermediate in the canonical pathway shown in A, while those that are unbolded produce non-canonical intermediates as shown in Fig. S4. N = 3 independent cultures; error bars = SD. (**C**), Representative micrographs of Pho8-GFP distribution in each strain. Only *erg3*Δ, *erg5*Δ, and *erg3*Δ*erg5*Δ and WT cells show robust domain formation in the vacuole, which is most abundant in the latter that synthesizes ergosterol. Scale bars, 5µm. Wider-field images with multiple cells are shown in Fig. S1A. (**D**), Quantification of domain formation frequency in fused vacuoles of stationary phase cells in 0.4% glucose. Mutants in bold predominantly produce an intermediate in the canonical pathway shown in A. Domains arise along the post-synthetic steps. N = 3 independent cultures; error bars = SEM.

Ergosterol’s post-synthetic steps are not essential for yeast viability, so combinations of gene knockouts could be used to test the activity of the preceding precursors (Fig. 3A). A metabolic map showing the products accumulating in each mutant strain is shown in Fig. S4. Among single gene knockouts, expected sterol intermediates were the most abundant sterol in *erg6Δ* (zymosterol) and *erg4Δ* (ergosta-5,7,22,24(28)-trienol), with minor accumulation side products or derivatives, as has previously been observed (*30*). The expected products of *erg2Δ* (fecosterol), *erg3Δ* (episterol), and *erg5Δ* (ergosta-5,7,24(28)-trienol) were modified by varying degrees to alternative, non-canonical sterols by subsequent enzymes in the pathways. To isolate pure canonical intermediates, we generated an *erg5Δerg4Δ* double knockout to isolate ergosta-5,7,24(28)-trienol, an *erg3Δerg5Δerg4Δ* triple knockout for episterol, and an *erg2Δerg3Δerg5Δerg4Δ* quadruple knockout for pure fecosterol. However, we continued analysis of non-canonical sterol-producing strains, since they provided additional information on structure-function relationships. Abundances of the canonical intermediates are shown in Fig. 3B and those for the non-canonical intermediates in Fig. S5.

Imaging of vacuoles of late-stage single knockouts showed that only *erg3Δ* and *erg5Δ*, which produce ergosta-7,22-dien-3β-ol and ergosta-5,7-dienol respectively, showed observable vacuole domains prevalent in WT cells (Fig. 3C). Among double knockouts, *erg3Δerg5Δ* cells accumulating ergosta-7-en-3β-ol also showed domain capacity, but only in a few cells. Cells synthesizing zymosterol (*erg6Δ*), fecosterol (*erg2Δ* and *erg2Δerg3Δerg5Δerg4Δ*), ergosta-7,22,24(28)-trien-3β-ol (*erg3Δerg4Δ*), and ergosta-7,22,24(28)-trienol (*erg5Δerg4Δ*) did not display any vacuole domains (Fig. 3D). Notably, all post-synthetic intermediates that supported vacuole domain formation lacked the C-24(28) alkyl tail unsaturation due to the activity of the C24(28) reductase Erg4. Many, but not all, late-stage mutants also showed a vacuole fragmentation phenotype as previously observed (*31*) (Fig. S1B), suggesting that ergosterol structural features are required for vacuole fusion that occurs when cells enter stationary phase (*32*). Enhanced fragmentation was also apparent in early-stage knockdowns to a lesser extent (Fig. S1C). Because fragmented vacuoles never showed membrane domains, vacuole fusion is likely required for domain formation. When accounting for cells containing successfully fused vacuoles, domain frequency in some mutant strains, like *erg5Δ*, approach but did not fully match those of WT cells (Fig. 3D). Vacuole domain frequency among all cells is provided in Fig. S1D.

As with early-stage mutants, we asked if observed phenotypes corresponded to domain formation capacity in model membranes. Since these intermediates are not synthetically available, we isolated sterol extracts from each strain (Fig. S6) and incorporated them as a 20 mol % sterol fraction into DOPC/DPPC (Fig. 4A). In the resulting GUVs, L_o_/L_d_ phase separation was supported by extracts from all strains that consistently displayed vacuole domains: *erg3Δ*, *erg5Δ*, and WT. We also observed phase separation in mixtures containing sterol extracts from *erg6Δ* (zymosterol) and *erg2Δ* (fecosterol) cells, but the resulting domains were irregularly shaped and did not coalesce during extended (> 1 hr) incubation at room temperature, indicating that they were solid or gel-like (*16*). Solid domains were previously observed in vesicles prepared with synthetic zymosterol, the primary sterol in *erg6Δ* extracts (*33*). No consistent phase separation was observed for the other canonical or non-canonical intermediates, though some GUVs prepared with *erg3Δerg5Δ* extracts (ergosta-7-en-3β-ol) showed solid domains. Overall, our analysis indicated that 1) post-synthetic intermediates preceding Erg2 activity, which contain an Δ^8^ B-ring, support the formation of solid domains 2) intermediates with a Δ^7^ B-ring but containing a C24(28) tail unsaturation (preceding Erg4 activity) do not support lipid demixing and 3) sterols containing both a Δ^7^ B-ring and a reduced C24(28) alkyl tail support L_o_/Ld membrane phase separation (Fig. 4B).

**Fig. 4.**
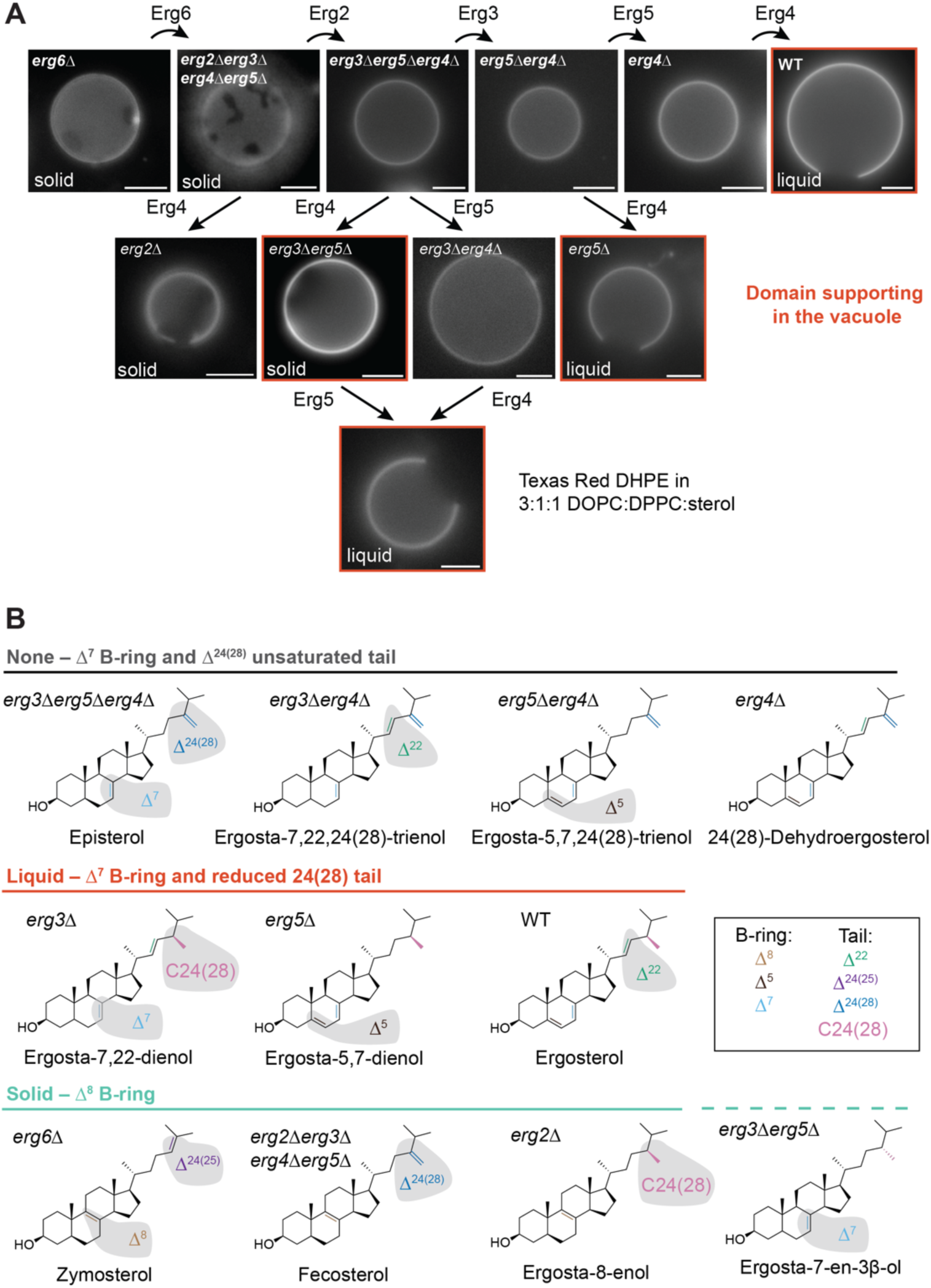
Structural features of ergosterol intermediates that support membrane domains. (**A**), Micrographs of GUVs composed of a 3:1 ratio of DOPC:DPPC with 20 mol % of sterols extracted from each post-synthetic mutant. GUVs contain the L_d_ marker TexasRed-DHPE and domains were allowed to coalesce at room temperature for >1 hr. Micrographs are arranged by their metabolic basis, with enzyme activity that converts their dominant sterol species shown through reaction arrows. In mixtures that support phase separation, the type is indicated by the inset text. Liquid domains are characterized by their capacity to coalesce. Extracts corresponding to yeast strains that show vacuole domains are highlighted in orange. The names of strains accumulating canonical intermediates are bolded. Scale bars, 5 µm. (**B**), Structures of the predominant sterols from each strain categorized by their ability to either promote gel-like domains (Solid), L_o_ domains (Liquid), or no domains (None) in GUVs. B-ring and tail structural features are color-coded to highlight differences in each category. Solid domain-promoting sterols contain a Δ^8^ B-ring, which is isomerized by Erg2, with the exception of ergosta-7-en-3β-ol that showed inconsistent solid domains. Sterols that cannot form domains under these conditions feature a Δ^7^ B-ring and have an unsaturated branched tail. Sterols that form liquid (L_o_) domains have a Δ^7^ B-ring and a tail that is reduced at C24(28) by Erg4.

### Non-monotonic changes in condensation capacity caused by post-synthetic modifications

The Bloch hypothesis predicts that membrane ordering of phospholipid membranes imparted by each sterol intermediate should increase along the pathway. However, ergosterol, as a final product, shows poor capacity to order unsaturated phospholipids when compared to cholesterol (*34*), suggesting that this model might not hold for the fungal pathway. To test this possibility, we used sterol extracts from the post-synthetic mutants and incorporated them into 1-palmiyoyl-2-oleoyl-PC (POPC) vesicles at a fixed stoichiometry (20 mol %) and assayed membrane ordering by the change in Generalized Polarization (ΔGP) of C-Laurdan compared to sterol-free POPC (Fig. 5A). In liposomes prepared from extracts of canonical intermediates, ΔGP was highest for early-stage intermediates (zymosterol, *erg6Δ*; fecosterol, *erg2Δerg3Δerg5Δerg4Δ*) and steadily decreased as the pathway progressed. Thus, intermediates that best support L_o_/L_d_ phase separation show low capacity to order membranes composed of the unsaturated phospholipids.

**Fig. 5.**
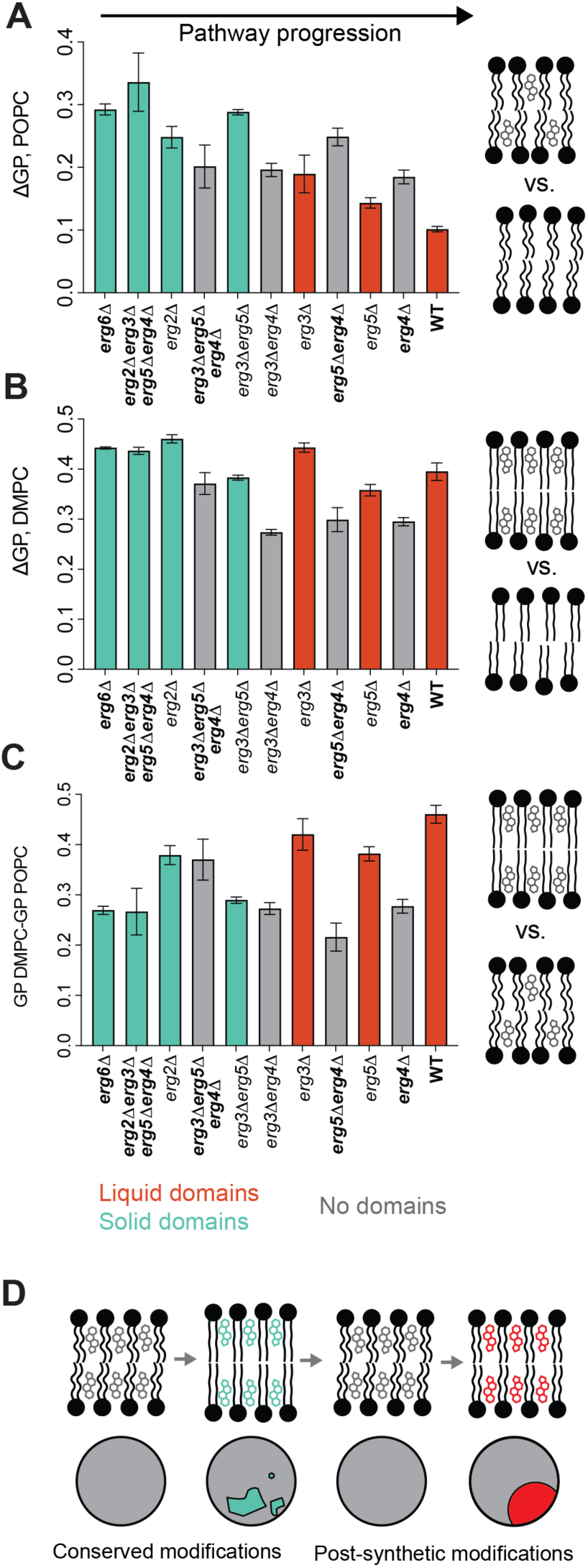
**Ordering capacities of ergosterol intermediates that support domains**. (**A**), The change in Laurdan GP for unsaturated (POPC) phospholipid vesicles upon incorporation of 20 mol % sterol was measured for extracts from each of the post-synthetic mutants. Mutants that accumulate canonical intermediates show bolded names. Strains whose sterols support membrane domains in GUVs are highlighted according to the type of domain. Liquid domain supporting sterols show low condensation of unsaturated phospholipids. N = 3 independent liposome preparations; error bars = SD. (**B**), The change in GP for saturated (DMPC) phospholipid vesicles upon incorporation of 20 mol % sterol. Domain supporting sterols show moderate to high ordering of saturated phospholipids, which corresponds to the melting temperatures of the mixtures (Fig. S7C). (**C**), The difference in GP between DMPC and POPC systems, both containing sterols, is largest for the ergosterol intermediates that also support liquid phase separation. (**D**), A model highlighting non-monotonic changes in ordering capacity in the ergosterol pathway and relationship to phase separation of coexisting domains of different types.

Given that ergosterol has been observed to have stronger ordering effects on saturated phospholipids (*34, 35*), we carried out identical experiments using saturated di-myristoyl PC (DMPC) mixtures vesicles (Fig. 5B). Like for POPC liposomes, we observed the highest ΔGP for early post-synthetic intermediates, consistent with previous measurements of zymosterol (*36*). In later steps, however, we also observed that ΔGP DMPC partially recovers for ergosterol compared to its immediate precursors, ergosta-5,7,24(28)-trienol (*erg5Δerg4Δ*) and 24(28)-dehydroergosterol (*erg4Δ*). Laurdan GP of DMPC with each sterol was correlated (R^2^ = 0.90) with the Tm of the mixtures, as measured using fluorescence polarization of di-phenyl hexatriene (DPH) (Fig. S7). When we calculated the difference in GP between POPC and DMPC liposomes containing sterol extracts, we observed ergosterol had the largest difference between the unsaturated and unsaturated systems (Fig. 5C). In general, extracts that allowed for L_o_/L_d_ phase separation in vacuoles and GUVs, such as *erg3Δ*, *erg5Δ*, and WT, also had large GP differences between DMPC and POPC systems. Thus, post-synthetic modifications do not uniformly increase ordering, but rather oscillate it maintain differences between sterol-containing unsaturated and saturated lipid membranes. This dynamic corresponds to the loss of solid domains and the reappearance of liquid domains along the pathway (Fig. 5D).

### The structural basis for sterol ordering explored by all-atom simulations

We were surprised by the capacity of small structural changes in ergosterol precursors – B-ring and alkyl chain unsaturations – to cause large changes in membrane ordering capacity. To further investigate the mechanisms underlying these effects, we employed molecular simulations of sterol-containing bilayers using the CHARMM36 all-atom force field. For each canonical post-synthetic intermediate, we generated and refined structural models, which were then incorporated into bilayers of di-myristoyl-phosphatidylcholine (DMPC) at 20 mol % (Fig. 6A). Using these simulations, we calculated changes in acyl chain ordering parameters (ΔS_CD_, averaged over acyl chain carbons four to ten) of DMPC chains relative to sterol-free systems (Fig. 6B). Overall, the correlation between ΔS_CD_ from simulations and melting temperatures measured experimentally for each extract in DMPC was robust (R^2^ = 0.92, Fig. 6C), supporting the use of CHARMM36 to interrogate sterol structure-function.

**Fig. 6.**
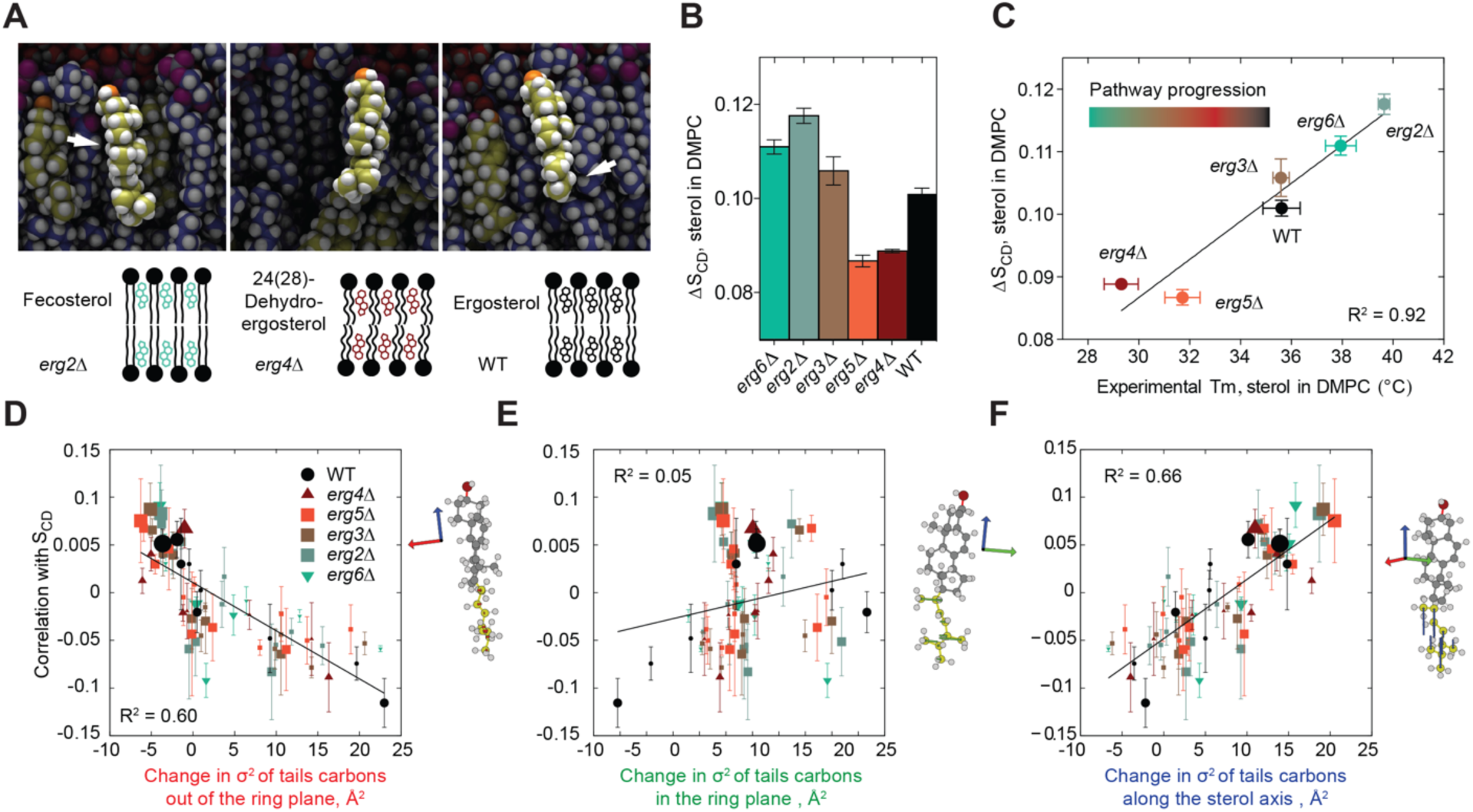
Atomistic models for phospholipid condensation by post-synthetic intermediates. (**A**), Examples of representative structures of post-synthetic sterol intermediates in all-atom simulations. The arrow in the fecosterol image highlights the smooth α-face of the ring system, while that in the ergosterol image highlights the extended alkyl tail. (**B**), Increase in acyl chain ordering parameter (ΔS_CD_) in simulations of DMPC with 20% canonical post-synthetic ergosterol intermediates compared to pure DMPC. Systems are referred to by gene deletion corresponding to the subsequent enzyme in the pathway that acts upon them. Experimental data for *erg2*Δ corresponds to fecosterol extracted from *erg2*Δ*erg3*Δ*erg5*Δ*erg4*Δ cells, *erg3*Δ corresponds to episterol from *erg3*Δ*erg5*Δ*erg4*Δ, and *erg5*Δ ergosta-5,7,24(28)-trienol *erg5*Δ*erg4*Δ. from Early intermediates show high ordering, which is then reduced, before partially recording in ergosterol (WT), as in experiments. Error bars = SD of 3 independent simulations. (**C**), ΔS_CD_ in simulations correlates with experimentally measured melting temperature (Tm) of the same mixtures (R^2^ = 0.915). Sterol-free DMPC has a Tm of 24 °C. Error bars = SD. (**D-F**), Correlation of S_CD_ with tail configuration shape relative to the sterol ring. For each dynamic sterol conformation, three dimensional axes (colored red, green, and blue) are determined by the moments of inertia of the ring system. Displacements of tail carbons are measured for each 3D axis, squared, and summed. The deviation of that sum from the mean is plotted on the x axis. The size of each mark represents their relative abundance for that specific sterol, which is indicated by its color and shape. Plotted are deviations out of the axis out of the ring plane (red, D), within the axis in the ring plane but perpendicular to the sterol long axis (green, E), and along the sterol long axis (blue, F). Small deviations out of the ring plane are positively correlated with order, as are extensions along the axis. There is no correlation for deviations within the ring plane. These features are consistent with an extended tail conformation correlating with ordering. Error bars = SD.

As in experiments, we found a significant difference in ordering capacity among post-synthetic intermediates. Early-stage sterols (zymosterol or *erg6Δ*, fecosterol or *erg2Δ*) showed high DMPC ordering capacities, mid-stage sterols (episterol or *erg5Δ*, 24(28)-dehydroergosterol or *erg4Δ*), showed low ordering capacity, and ergosterol displayed an intermediate level (Fig. 6B). This non-monotonic trend mirrored what was observed in membrane ordering experiments (Fig. 5). Visualization of these systems showed clear differences in the sterol conformations, particularly in the alkyl tail that is remodeled during the post-synthetic steps (Fig. 6A). To quantify this relationship, we correlated each sterol conformation, measured via each dihedral angle of the sterol tail, to the instantaneous total bilayer order in each frame of the simulations. This analysis showed that ordering was correlated with low deviations of the tail carbons out of the plane of the sterol ring (Fig. 6D). In contrast, there was no correlation between ordering and with variations along the ring plane (Fig. 6E). Ordering was also correlated with increasing extension of carbons along the sterol axis itself (Fig. 6F). Taken together, these results indicate that increased sterol ordering results from a more extended tail conformation in bilayers.

### Long-range organization of phospholipids modulated by ergosterol metabolism

To further understand the effect of differing sterol structures on neighboring phospholipids, we analyzed the distribution of DMPC ordering (S_CD_ parameters) across the entire bilayer for each sterol system. Across all systems, the distribution of individual acyl chain S_CD_ values could be deconvolved into two peaks (Fig. S8A), a tightly distributed high order population, and a broader, lower order one. The former increases with the average order of the system, while the latter is largely unchanged (Fig. 7A). Surprisingly, the differences in S_CD_ across all systems was explained entirely by the increase in frequency of the high-order configurations, as the abundance of these DMPC states was tightly correlated (R^2^ > 0.99) with total simulated membrane ordering (Fig. 7B) and experimentally measured melting temperatures (Fig. S8B). When we classified DMPC acyl chains by their *gauche*-*trans* isomerization, which yielded 1,024 distinct classes of chain conformers, we found that only ten chain classes present a monotonic percentage increase with increasing order. These classes all present at least 7 consecutive carbons with *trans* conformations: XTTTTTTTTX, TGTTTTTTTX, TTTTTTTTGX and TTTTTTTGTG, with X being either *gauche* (G) or *trans* (T). Thus, total membrane order arises through a distinct population of saturated phospholipids with mostly *trans* acyl chain conformations (Fig. S8C). The frequency of this population oscillates during successive sterol modifications in the post-synthetic steps.

**Fig. 7.**
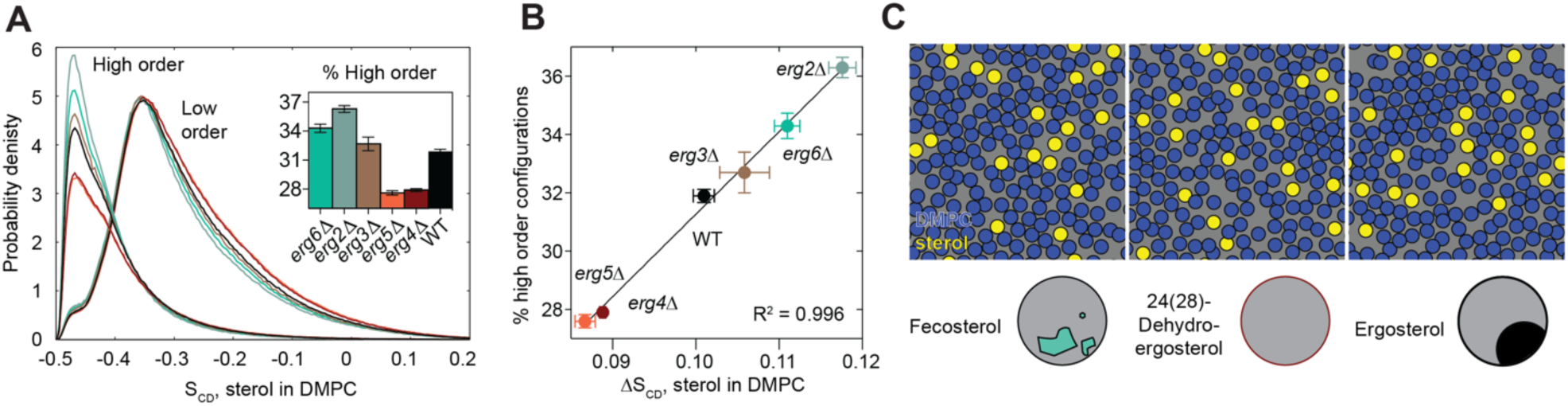
Post-synthetic intermediates modulate long-range ordering of saturated phospholipids. (**A**), The frequency of the high-order DMPC populations, which contains mostly *trans* acyl chain conformers, increases in abundance with higher ordering sterols. Error bars and total distributions for each system and sterol-free DMPC are shown in Fig. S8A. (**B**), The increase in higher order configurations tightly correlates with total ordering for each sterol system. Error bars = SD. (**C**), Sterol structural changes manifest in long-range ordering of DMPC. Panels show top-down view of the all-atom simulation with DMPC acyl chains and ergosterol rendered as blue and yellow circles, respectively. Transient pockets of ordered DMPC acyl chains are observed in all systems but are most extensive in the solid-domain forming fecosterol (left) and least extensive in sterols that do not allow for demixing, like 24(28)-dehydroergosterol (middle). Ergosterol (right), which supports fluid domains, shows an intermediate level.

One model for sterols ordering phospholipids is strictly through direct sterol-acyl chain interactions, which were envisioned initially by Bloch. However, in our simulations DMPC chains outnumber sterols 8:1, so the frequency of induced ordered chains we observed was in stoichiometric excess of the sterol (Fig. 7A, inset). Acyl chains near the smooth, α-face of the simulated sterols were also not more highly ordered in simulations compared to the overall average. Thus, the effect of changing sterol structure manifests in long-range ordering within the system. An explanation of this paradox lies in the assembly of transient, hexagonally packed regions of saturated acyl chains, which we observed as the 1 μs simulations progressed (Fig. 7C). Such packing was previously observed in simulations of liquid-ordered regions of ternary lipid mixtures that phase separate (*37, 38*). High-ordering sterols in the pathway, like fecosterol, led to systems with widespread hexagonal packing DMPC, while systems with no or only low-ordering sterols, like 24(28)-dehydroergosterol, had few such structures. Ergosterol systems showed an intermediate level, with regions of dense hexagonal packing and other regions of disorder. This feature corresponds with the sterol’s capacity to support ordered domains that remain fluid.

## Discussion

Here we explored the biophysical driving forces underlying a single metabolic pathway through a combination of *in vivo*, *in vitro*, and *in silico* approaches. Previous comparisons between sterols have largely focused on a small number of natural or synthetic sterols. We capitalized on the ability of yeast cells to remain viable while accumulating intermediates across the ergosterol pathway to perform a systematic analysis. The resulting strains (Table S1) are broadly useful for systematic investigation of sterol function. Here, we used domain formation of the yeast vacuole as a cellular readout of a sterol-dependent property (membrane phase separation), generating structure-predictions that could be tested by model membrane experiments and through simulations. These additional approaches are important because our analysis of sterol composition *in vivo* did not measure their abundance in the vacuole membrane, which is controlled by trafficking, nor account for potentially compensatory changes in other lipid components. These might explain why some behaviors observed in synthetic vesicles, like gel domains caused by early post-synthetic intermediates, were not observed in vacuoles. Nonetheless, we found agreement between several observations made in vacuoles and the propensity of extracted intermediates to support domain formation in vesicles, and all sterols that supported regular domains in the former yielded L_o_/L_d_ phase separation in the latter. We also used sterol extracts from mutants to characterize the ordering capacity of post-synthetic intermediates, which agreed with all-atom simulations based on sterol models built with permutations from the metabolic pathway. Analysis of simulations showed that unsaturations in the sterol B-ring and alkyl tail are potent modulators of ordering capacity, which manifests through nanometer-scale hexagon packing of saturated chains, as previously observed in simulated liquid ordered phases (*37, 38*). That is, total order is a measure of the sterol’s ability to promote the collective structure of the ordered phase, rather than acting completely locally.

Our experimental design was motivated by the Bloch hypothesis, which postulated that the evolution of long metabolic pathways for sterols would have required a progression of biophysical fitness along the pathway. We find that the ergosterol pathway matches the criteria laid out by Bloch, but not for phospholipid condensation as originally proposed. Instead, the pathway acts to generate a final product that can interact with phospholipids in a manner that supports phase separation of coexisting fluid domains. Counterintuitively, it does so by reducing the condensation capacity of its final product, ergosterol. Initial demethylated sterols, like zymosterol and fecosterol, provide too high of an ordering capacity, and drive the formation of solid domains in model membranes. In simulations, these intermediates show a smooth-faced ring system (Fig. 6A), cause a high abundance of phospholipids with all *trans* acyl chain isomers (Fig. 7B), and promote extensive hexagonal packing that is consistent with the nucleation of solid domains (Fig. 7C). The pathway subsequently acts to reduce condensation capacity through the Δ^8^ isomerase Erg2 and further through two additional desaturases acting at the B-ring (Erg3) and alkyl tail (Erg5). This generates a series of intermediaries with poor ordering capacity and an inability to support any phase separation. These low-ordering sterols, like penultimate intermediate 24(28)-dehydroergosterol, feature a rougher ring system and promote only low levels of long-range ordering compared to earlier intermediates. The final step of the pathway, carried out by the reductase Erg4, generates a saturated methyl group at the alkyl tail, which partially recovers ordering capacity in ergosterol and the two other intermediates (ergosta-5,7-dienol and ergosta-7,22-dienol) that support phase separation in both vacuoles and GUVs. These Erg4 products still show low ordering capacity for unsaturated lipids, but regain an intermediate ordering capacity for saturated lipids (Fig. 5 and Fig. 6B).

Compared to all its precursors, we find that ergosterol optimizes the phase separation of membranes into coexisting liquid phases. Previously, it was assumed that formation of L_o_ domains correlated with a sterol’s capacity to order phospholipids (*20*). However, the capacity of ergosterol to form ordered domains, despite its modest ordering capacity, has been unexplained (*39*). In yeast cells, phase separation of membrane domains is required for cell survival, as it underlies micro-autophagy in the vacuole (*23, 40*). Similar processes have been proposed to act in numerous biological processes in other cell types (*41, 42*). Unlike condensation capacity, formation of liquid domains depends on sterols and their interactions with at least two other classes of lipids, e.g. unsaturated and saturated phospholipids with low and high melting temperatures (*16*). Membrane phase separation requires sufficient line tension between two coexisting phases (*43, 44*), which is related to their thickness differences (*45–47*). Consistent with these models, we observe that ergosterol still supports a large ordering difference between unsaturated (POPC) and saturated (DMPC) phospholipids. A challenge for biologically relevant membrane phase separation, however, is achieving such ordering differences while preventing domains from falling below their melting temperature. While ergosterol’s moderate ordering capacity supports liquid phase separation, early demethylated precursors (zymosterol and fecosterol) that induce higher ordering instead drive formation of solid domains. They do so by causing too high of an abundance of phospholipids with all *trans* acyl chain conformers. This dynamic could explain why ordering capacity must be tuned down by post-synthetic steps in the ergosterol pathway.

As Bloch first proposed, the properties of sterol intermediates carry implications for the evolution of sterol metabolism in eukaryotes. In the initial steps of sterol metabolism, an ability of linear precursors, squalene and 2,3-oxidosqualene, to promote membrane heterogeneity (Fig. 2C, 2E) suggests that they could have served early roles in membrane organization. The synthesis of these terpenes and their cyclization into lanosterol could have acted to promote or reduce gel-like domains, respectively, alongside sphingolipids. Conserved modifications, which demethylate lanosterol into zymosterol, would have been selected for in the last common ancestor of eukaryotes due to the elevated membrane stability and reduced permeability provided by demethylated sterols, as postulated by Bloch. The high ordering capacity of early ergosterol precursors, like fecosterol, could also explain why their accumulation has been selected for in laboratory evolution experiments at high temperatures that otherwise reduces membrane ordering (*48*). In the post-synthetic steps, a requirement for functional membrane organization would have driven the multi-step adoption of ergosterol, which has reduced ordering capacity compared to zymosterol but allows for L_o_/L_d_ phase separation. Within the post-synthetic steps, the promiscuity of Erg4’s C24(28) reductase activity and its capacity to support domains in both *erg3Δ* and *erg5Δ* backgrounds suggests that it might have evolved prior to Erg3 and Erg5’s desaturase activity. In this way, fluid membrane domains would have arisen before the adoption of ergosterol as a final product, even if the direct precursors of ergosterol in the extant pathway do not support them. In this putative proto-pathway (a sequence of Erg6–Erg2–Erg4–Erg3–Erg5), successive final products during the pathway’s evolution would have initially supported solid domains, followed by a series of increasingly robust fluid domains.

While ergosterol is the dominant sterol across fungi, metazoan and plant sterols can also support phase separation both in model systems and in cells. The biosynthetic pathway(s) for cholesterol, which Bloch first characterized, shows intriguing similarities and differences from that of ergosterol. Both feature a Δ^8^ isomerase step that buffers the capacity of their shared early intermediates to form solid domains and late-pathway reductases that are essential for generating a product that supports phase separation (*19*). However, C24 methylation, found in ergosterol and some plant sterols, is absent from cholesterol metabolism, and so these late-stage reductases act on different positions, e.g. C24(25) and at the B-ring. How these different pathways diverged remains an open question, but their evolution suggests multiple chemical solutions to achieve membrane organization. Notably, enzymatic promiscuity is common in sterol metabolism; switching between two sterols can even occur via single step changes much earlier in pathways (*49*). Generation of new sterols could have occurred rapidly during early eukaryotic evolution, potentially in response to the diversification of other membrane components, like sphingolipids whose chemistry also varies across eukaryotic lineages (*12*). In this way, the radiation of sterol metabolism could have also been driven by the need to balance interactions with other lipids.

## Materials and Methods

### Yeast strains, plasmids and media

*S. cerevisiae* W303a was the base strain for the study. Knockout strains were generated by PCR-based homologous recombination. For knockdown strains except those targeting *ERG7*, a tetO2-*CYC1* promoter and expression cassette for the tetracycline-controlled transactivator were amplified from plasmid PCM224 and substituted upstream from the start codon of the gene of interest. For *ERG7* promoter substitution, the *MET3* promoter was utilized instead. All strains generated are listed in Table S1. For visualization of vacuole domains *in vivo*, yeast strains were transformed with plasmid pRS426 GFP-Pho8. For strain generation, cells were grown in YPD medium (Fisher Scientific), buffered complete supplement mixture (CSM, 0.5% ammonium sulfate, 0.17% yeast nitrogen base without amino acids, 80 mM potassium phosphate dibasic and 2% glucose), and buffered minimal medium (0.5% ammonium sulfate, 0.17% yeast nitrogen base without amino acids, 80 mM potassium phosphate dibasic and 0.4% glucose).

### Analysis of vacuole membrane domains

Vacuole membrane domain formation was induced by glucose depletion and cells were subsequently imaged by confocal microscopy, as described previously (*50*). Briefly, a starter culture harboring the pRS426 GFP-Pho8 plasmid is grown in 5 mL YPD overnight then diluted 1:100 into 5 mL CSM medium lacking uracil for selection and incubated for 16-18 hrs. Approximately 0.1 OD unit of the CSM culture was diluted in 5 mL of minimal medium containing 0.4% (w/v) glucose and lacking uracil and incubated for 24 hrs prior to imagining. For *ERG* knockdowns under control of the tetO2-*CYC1* promoter, a 10 mg/mL stock solution of doxycycline (Sigma-Aldrich) was added to the minimal medium to a final concentration between 100 ng/mL to 500 ng/mL. For *ERG7* a 1000X (20 g/mL) methionine stock solution (Sigma-Aldrich) was added to the minimal medium at a final concentration of 2X (40 mg/mL). Cells growth in minimal medium were transferred to 8-well microscope chamber slides (Nunc Lab-Tek) pre-coated with 1-2 mg/mL concanavalin-A (MP Biomedicals). Vacuoles were imaged using a Zeiss LSM 880 Airyscan microscope at room temperature with a Plan-Apochromat 63x/1.4 Oil DIC M27 objective. GFP excitation was via a 488 nm Argon laser at 2% power. Images were processed in Zen Black using default Airyscan settings. Processed images were categorized into groups based on vacuole domain morphology and vacuole fragmentation using the multi-point tool in ImageJ as described previously (*50*). Vacuoles were captured from top to bottom and 3D projections were generated using the Z project tool in ImageJ. A minimum of 100 cells were quantified in each biological replicate.

### Cellular sterol extraction and analysis

Extraction of sterols from yeast has been described previously (*51*). Briefly, an amount of cells corresponding to an absorbance of 1 at OD600 was spiked with internal standard (cholesterol) and resuspended in 1 mL of sterile MQ water. 1 mL of glass beads was added to break apart the cells, the resulting cell lysate was then transferred to a 15 mL conical glass with a teflon cup. 3 mL of methanolic KOH was added. The resulting mixture was incubated at 70 °C for 2 hours. The mixture is allowed to cool and 3 mL of *n*-hexane was added. The mixture was then mixed with a vortex at the highest setting for 10 seconds. The tube was then spun at RT by centrifugation for 5 minutes at 800 RPM. The upper organic phase was then transferred to a clean conical glass tube. A second extraction of the aqueous phase was performed and the resulting organic phase was combined. The n-hexane was evaporated under a stream of nitrogen. The resulting dry extract was then dissolved in a 1:1 solution of pyridine and TMCS and incubated at 70 °C for one hour for derivatization. Extracts were analyzed by GC-MS as follows. Samples were injected into a 30m DB-5 column in an Agilent 8890 GC coupled to a 5977B mass analyzer. The GC oven was operated with the following temperature program: initial temperature 120 °C held for 1 min, ramped at 20 °C per min to 270 °C, followed by an additional ramp at 2 °C per min to 290 °C, then a final ramp at 20 °C per min to 300°C and held for 2 min. The identities of target compounds were initially checked against the NIST17 mass spectral library using NIST MS Search software. Mass spectra from the literature were used as an additional source when target compounds could not be found in the library. For quantification, sterol levels were calculated relative to a calibration curve of a cholesterol external standard and adjusted according to the extraction efficiency for each sample. Extraction efficiency was determined on a percent recovery basis of internal standard per sample.

### Generation and analysis of sterol extracts

A starter culture in 5 mL CSM was prepared and grown in a shaker for 24 hrs, the culture was diluted 1/100 into a final volume of 50 mL with fresh CSM in a 250 mL flask. After 24 hrs incubation the culture was diluted 1/100 into a final volume of 650 mL with fresh CSM in a 2L non-baffled flask shaking at 200 RPM for 24 hrs. Cells were harvested by benchtop centrifugation (3000xg for 20 minutes). Cells were washed with 650 mL of MQ water, then resuspended again in 650 mL MQ and aliquoted into 50 mL falcon tubes. Unused, cells were spun down again (3000 RPM for 10 minutes) and MQ water removed and stored in the -80C for future use. For cells that were to be used immediately 5 mL of resuspended cells were transferred into a 15 mL falcon tube for sterol extraction as described above with modifications. After alkaline hydrolysis, the unsaponified fraction was extracted with hexane. An additional cleanup step was performed to reduce the amount of non-sterol lipids present in the unsaponifiable fraction as described elsewhere (52). Briefly, the unsaponified fraction was mixed in a solvent mixture of additional hexane, acetone, and methanol such that the composition was 72% (v/v) hexane, 14% (v/v) acetone, 14% (v/v) methanol. This mixture was extracted in a separatory funnel with a solvent mixture consisting of 39.6% (v/v) hexane, 17.82% acetone, 41.58% methanol and 1% (v/v) water. After separation the upper organic phase was concentrated by rotary evaporation. The resulting lipid film was resuspended in 2 mL of chloroform and stored in an airtight container (Avanti 600511) at -20 °C until use. Extracts were quantified by GC-MS, as described above, and quantified using an ergosterol external standard. GUVs and liposomes from sterol extracts were prepared as described above.

### Giant vesicle electroformation

Phosphatidylcholine (PC) (Avanti Polar Lipids), ergosterol (Thermo Scientific), squalene (Echelon Biosciences), squalene (Sigma-Aldrich), 2,3-oxidosqualene (Sigma-Aldrich), GluCer C16 (Avanti Polar Lipids), lanosterol (Avanti Polar Lipids), FF-MAS (Avanti Polar Lipids), and Texas Red dihexadecanoyl-PE (DHPE; Life Technologies) were used as purchased without further purification. Lipid solutions in chloroform typically contained dipalmitoyl-phosphatidylcholine (DPPC, 16:0 PC), ergosterol, 0.8 mol % Texas Red DHPE, a dye that selectively partitions to the L_d_ phase. GUVs were electroformed in 200 mM sucrose, 0.25 mg of lipids in stock solutions were spread evenly on slides coated with indium tin oxide (Delta Technologies). Lipid coated slides were placed under vacuum for > 30 minutes to evaporate the chloroform. A capacitor was created by sandwiching an O-ring (Ace Glass 7855-14) between two lipid-coated slides, filling the gap with sucrose solution. An AC voltage of 10 Hz and 1.3 V was applied across the capacitor for 1 hr at 60 °C. GUVs were harvested and allowed to cool at room temperature for 1 hr. Phase separation was assessed by imagining under wide-field fluorescence microscopy (Thermo EVOS equipped with 63x Nikon oil immersion objective).

### Fluorescence spectroscopy

Liposomes for spectroscopy were generated by thin film hydration at 50°C and extruded 21 times through 100 nm filters (Avanti Polar Lipids). Steady state fluorescence readings were performed on a Cary Eclipse fluorometer (Agilent) equipped with automated polarizers. GP of Laurdan (Invitrogen D250) was calculated from emission intensities at 440 nm and 490 nm upon excitation at 365 nm in liposome suspensions extruded to 100 nm and stained with 50 µM of the dye. Anisotropy of DPH (Sigma D20800) was calculated from emission at 430 nm upon excitation at 360 nm in liposomes stained with 50 µM of the dye. All measurements were taken at 30 °C. For estimation of Tm, DPH anisotropy values were obtained across a temperature range (T) of 8°C to 70°C and fitted to a Boltzmann sigmoidal equation:

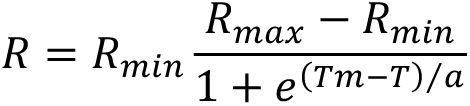

Where R refers to the DPH anisotropy ratio, and R_min_ and R_max_ are the minimum and maximum fluorescence, respectively, T is the temperature, Tm is the melting temperature, and a is the fitted slope.

### Molecular dynamics simulations

Except for ergosterol itself, simulated ergosterol precursors models are named by the genetic change that would result in its upregulation, assuming the pathway in Fig. 3A, as opposed to their chemical name. Systems were built using the CHARMM-GUI web module with ergosterol precursors substituted for ergosterol as needed. Parameters and chargers were chosen by analogy to similar chemical structures in the CHARMM C36 lipid forcefield. In the one case where parameters were not obviously transferrable (the diene torsion of 24(28)-dehydroergosterol in *erg4*Δ models) dihedral parameters were fit to a model compound (the tail of 24(28)-dehydroergosterol) using the MP2/6-31G* quantum chemical method and the Q-Chem 5.0.1 software package. Each system was initially equilibrated using the standard steps of the CHARMM-GUI package using the molecular dynamics package NAMD. Subsequent equilibration and production were performed with AMBER 20 on a single A100 GPU. Standard simulation parameters were used appropriate for the C36 forcefield (Lennard-Jones forces were switched off between 10 and 12 Å, the particle mesh Ewald method was used to compute long range electrostatics, bonds to hydrogen were constrained using SHAKE/SETTLE). Temperature was maintained at 30 °C using a Langevin thermostat. Isotropic 1 atm pressure at zero lateral tension was maintained using AMBER’s Monte Carlo barostat. Systems were run at 20% sterol in DMPC (20 sterol molecules and 80 phospholipids per leaflet) for at least 500 nanoseconds each, in triplicate, with the initial 100 nanoseconds neglected. Approximately 3700 water molecules were used per system, consistent with the default hydration recommended by CHARMM-GUI.

### Sterol tail conformation analysis

Tail conformations were identified by first constructing histograms of each independent dihedral angle of a sterol tail. Stable dihedral basins with significant likelihood were unambiguous. Each basin of each dihedral was assigned a discrete number, enabling enumeration of all possible tail conformational states. For each frame of simulation analysis, the instantaneous total bilayer order was correlated with the instantaneous number of each tail conformation. For example, the most likely tail conformation of ergosterol [44%; C13-C17-C20-C22 176°, C17-C20-C22-C23 239°, C22-C23-C24-C25 117°, C23-C24-C25-C26 291°] had correlation ρ=0.056 with the order parameter. The impact on overall order ΔS_CD_ from conformation *i* was estimated as:

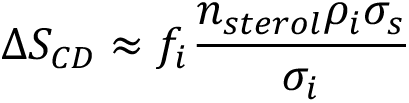

where σ_s_ is the standard deviation of the order parameter, *f*_i_ is the average fraction of sterol with conformation *i*, σ_i_ is the standard deviation of *n*_sterol_*f*_i_, and ρ_i_ is the correlation of bilayer order with the number (*n*_sterol_*f*_i_) of conformations *i*. This is equivalent to a least-squares linear fit to the order as a function of the total number of tail conformations, evaluated at the observed number of tail conformations.

### Acyl chain population analysis

For all the systems, the ten dihedral angles from the DMPC acyl chains (carbons 2 to 14) were computed. Dihedrals with absolute angles smaller than 115° were considered *gauche* (G), others *trans* (T). Each acyl chain was then assigned a ten-letter code with either G or T in each position. The percentage of each code was computed for all systems, revealing a monotonic increase of codes XTTTTTTTTX, TGTTTTTTTX, TTTTTTTTGX and TTTTTTTGTG (total of 10) with increasing order. The distribution of order parameters (S_CD_) for those configurations was then generated for each system, showing a significant contribution to the high order peak of DMPC acyl chains, which again increased with order (Fig. S8).

## Acknowledgements

Edward Lyman, and Sarah Keller provided discussions that informed the work. Jacob Winnikoff provided helpful comments on the manuscript. Funding was provided by the National Science Foundation (MCB-2046303 to I.B.), the Department of Energy (DE-SC0022954 to I.B.), and the National Institutes of Health (NIH) (1ZIAHD008955 to A.S., GM142960 to I.B., T32-GM008326C to I.J-C.). L.J.S.L. and A.J.S. were supported by the intramural research program of the National Institute of Child Health and Human Development (NICHD). Computational resources were provided by the NICHD and by the NIH HPC Biowulf cluster.

---

**Fig. S1.**
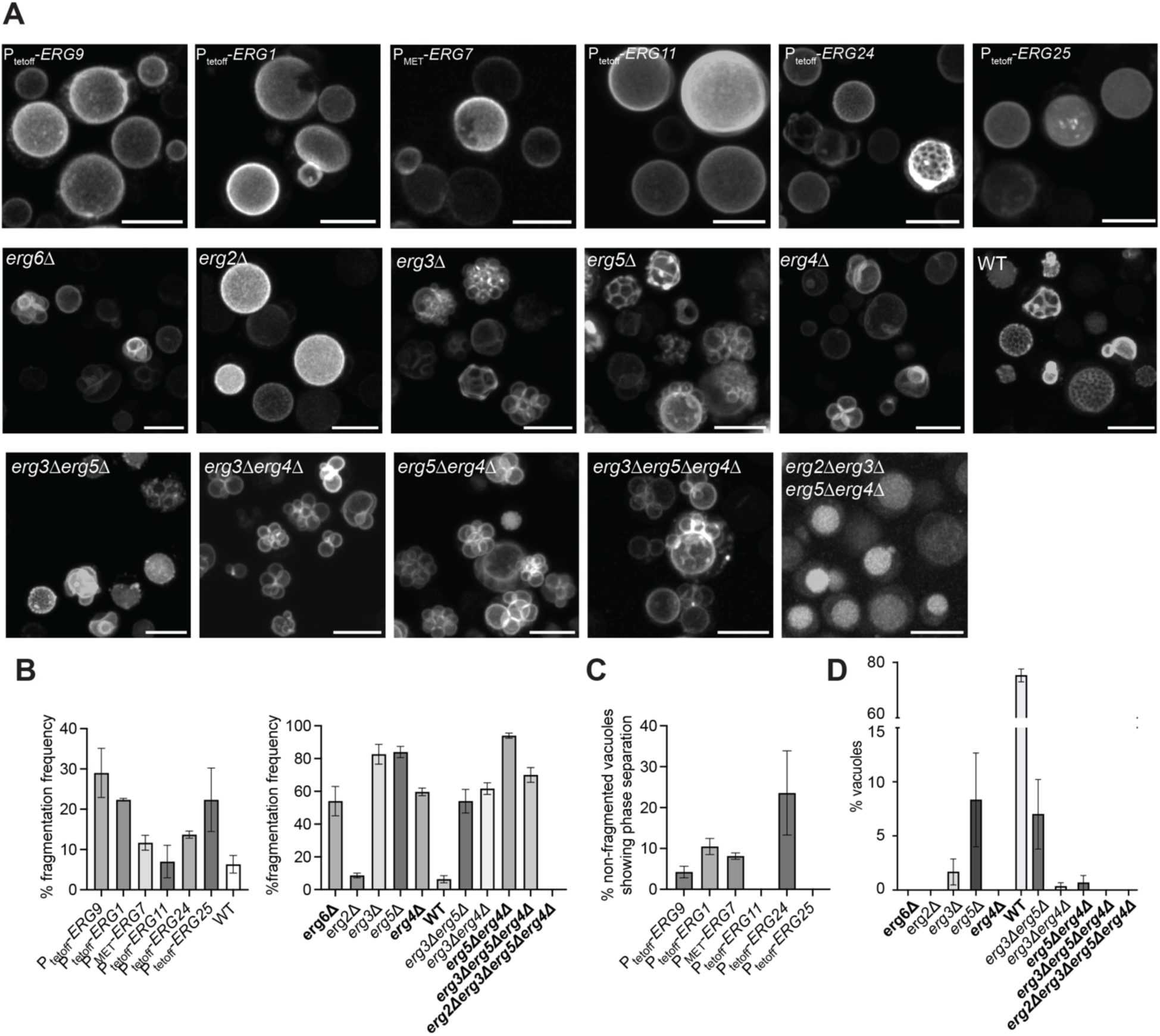
Additional characterization of vacuoles in ergosterol mutants. (**A),** Representative micrographs for Pho8-GFP and vacuole morphology distribution in fields of cells for each strain. Scale bars, 5µm. (**B**), Quantification of vacuole fragmentation frequency across strains, which is sterol dependent. Only *erg2Δ*, and WT cells show <5% vacuole fragmentation, whereas *erg2Δerg3Δerg5Δerg3Δ* exhibits no fragmentation. Post-synthetic mutants in bold indicate strains that accumulate the canonical ergosterol intermediates. (**C**), Quantification of early-stage mutant domain formation frequency considering only fused vacuoles. Fragmented vacuoles universally did not exhibit domain formation under any conditions. (**D**), Quantification of post-synthetic mutant domain formation frequency considering all vacuoles, including fragmented ones that did not undergo fusion when entering stationary phase.

**Fig. S2.**
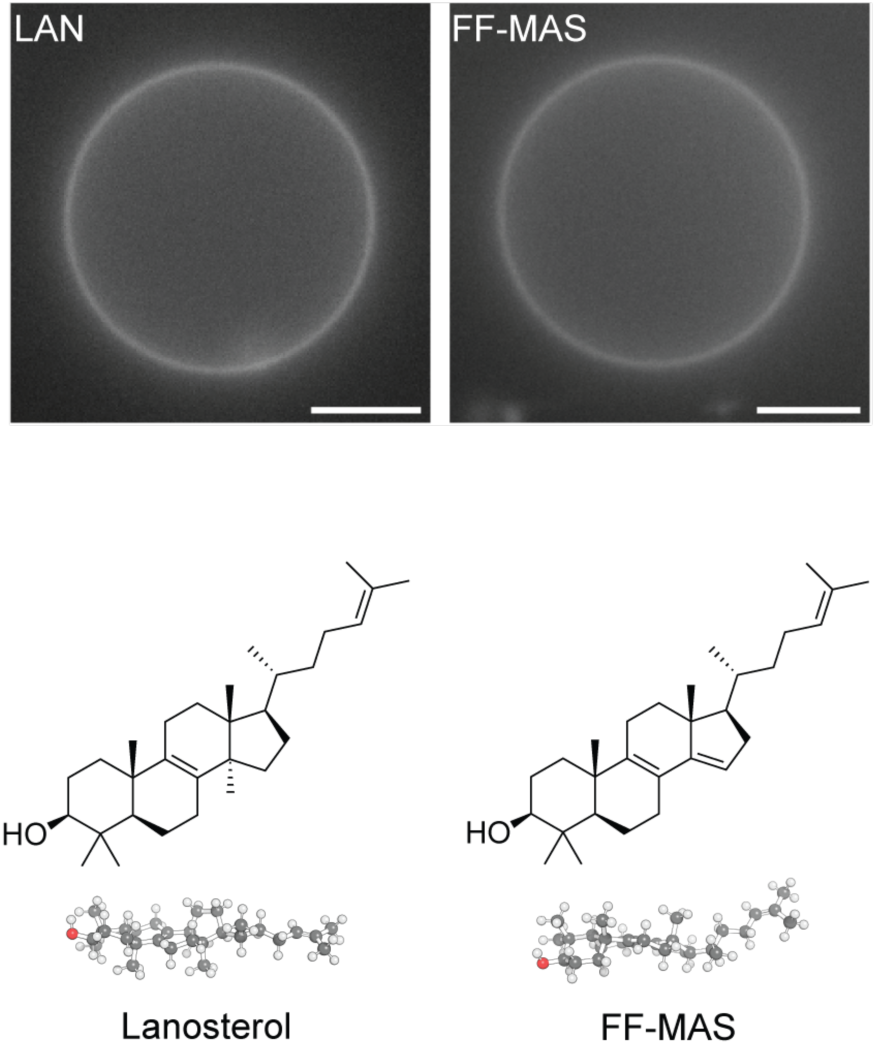
Methylated sterols including lanosterols (LAN) and FF-MAS do not support domain formation. Images show GUVs containing 3:1 DOPC:DPPC with 20 mol % of sterol. Ld regions are labeled with TexasRed DHPE. Scale bars, 5 µm.

**Fig. S3.**
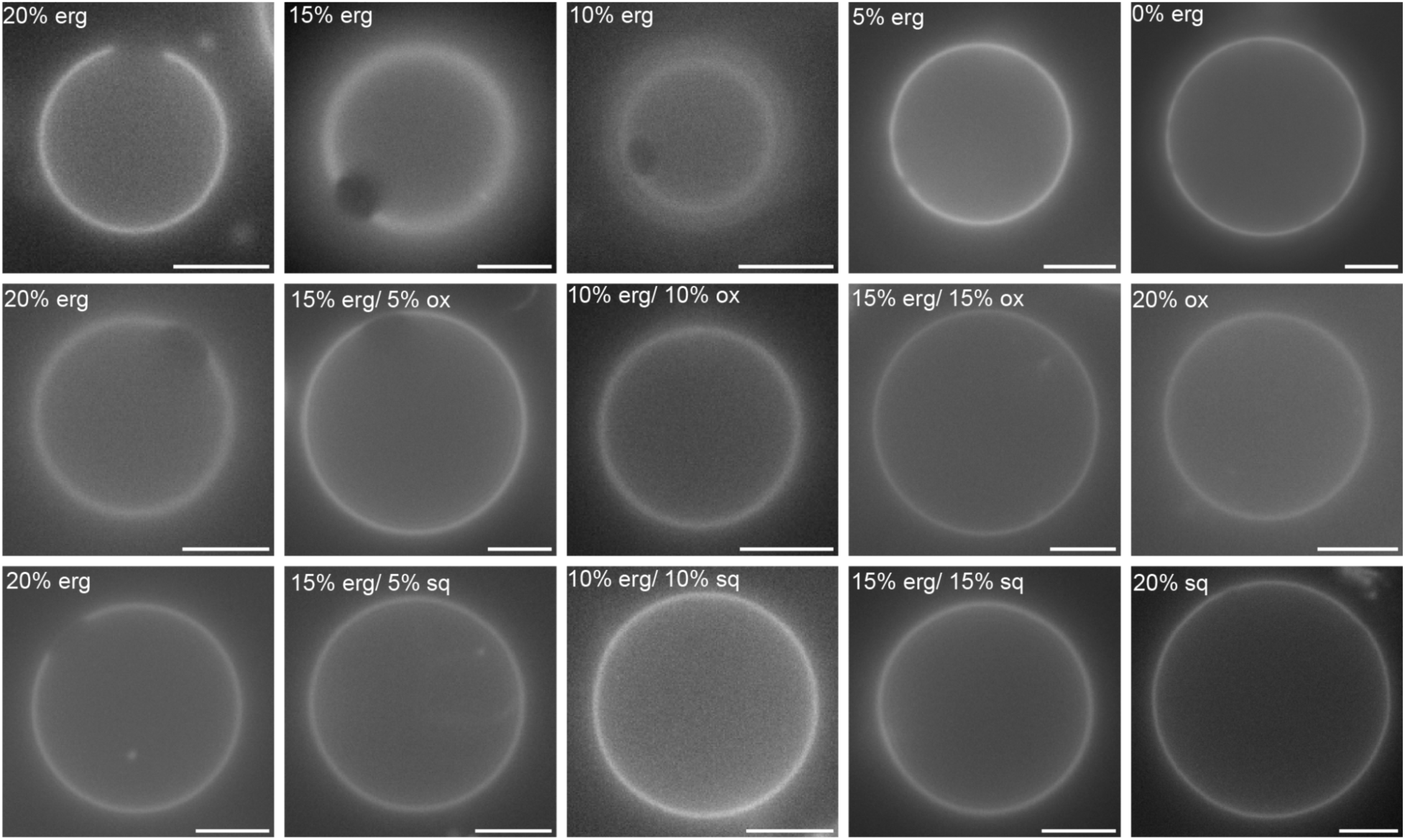
Squalene (sq) and 2,3-oxidosqualene (ox) do not support L_o_/L_d_ domains in the absence of sphingolipids. Top Row: GUVs composed of a 3:1 ratio of DOPC/DPPC with varying ergosterol compositions suggest a minimum ergosterol threshold of 10% required for domain formation to occur. Middle Row: GUVs composed of the same DOPC:DPPC ratio with decreasing amounts of ergosterol supplemented with increasing 2,3-oxidosqualene show an increased threshold of 15% ergosterol required to support domains when supplemented with 2,3-oxidosqualene. Bottom Row: Similarly, squalene accumulation results in a minimum ergosterol threshold of 20% required for microdomain formation to occur.

**Fig. S4.**
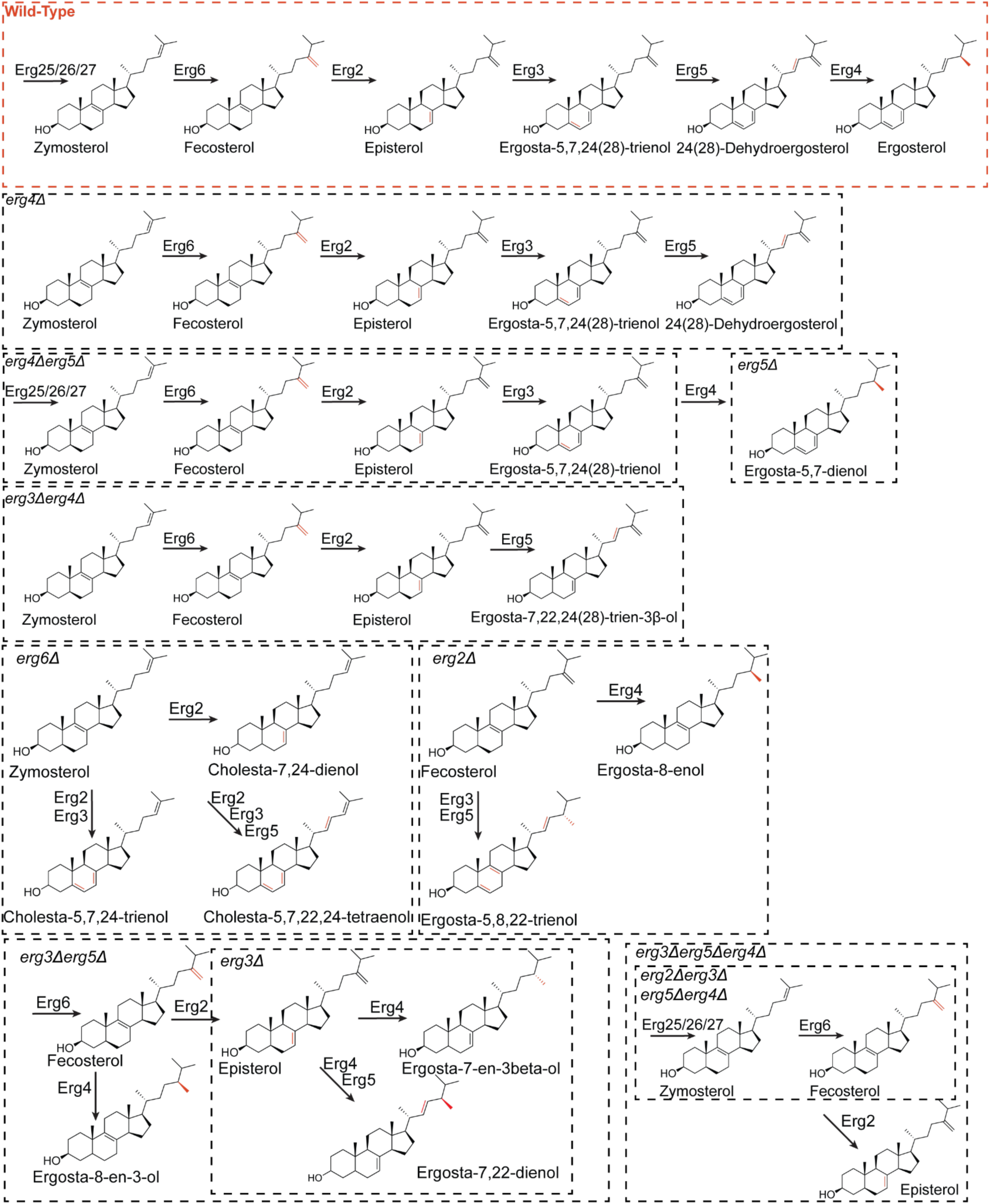
Metabolic map showing sterols products across all post-synthetic pathway mutants. Dashed boxes indicate the pathways relevant in the primary product of each strain.

**Fig. S5.**
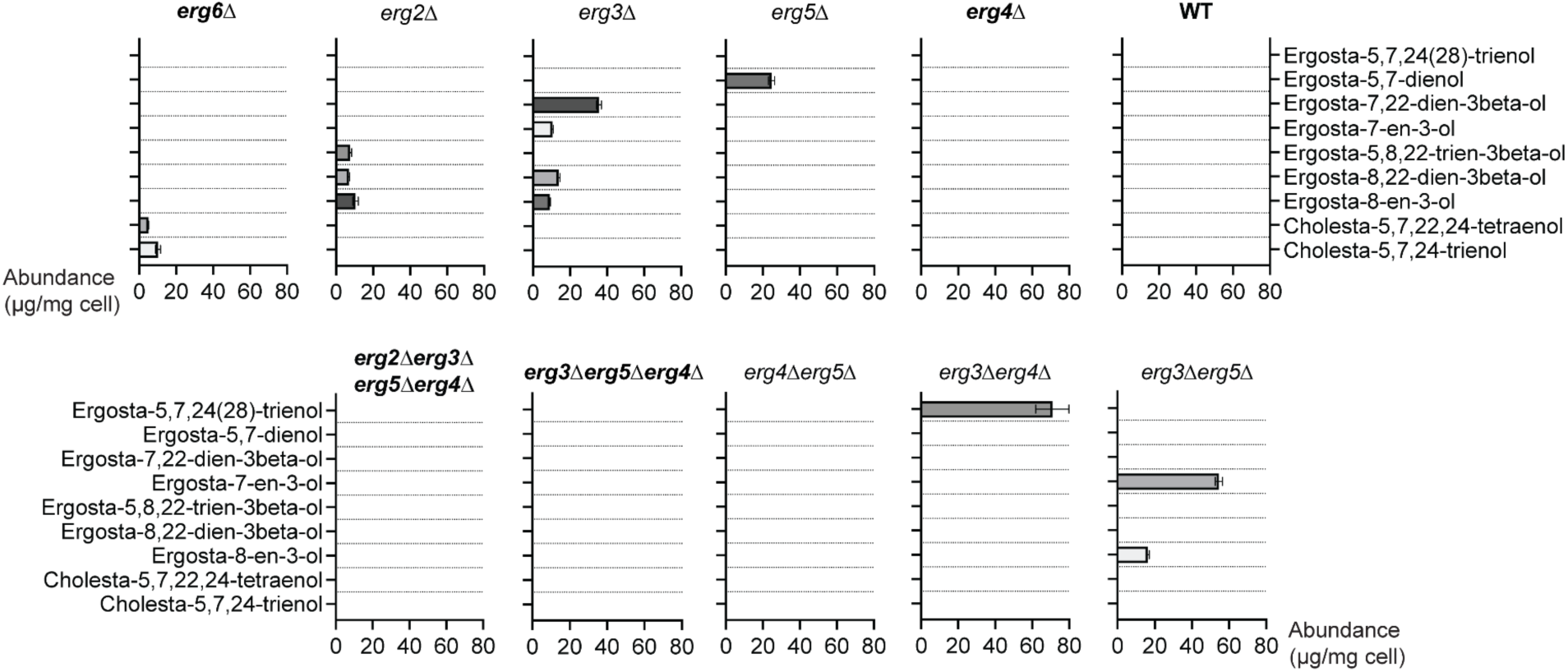
Abundances of non-canonical ergosterol intermediates in each post-synthetic mutant strain, as measured by GC-MS. Bolded strain names predominantly accumulate single, canonical intermediates. Structures for each intermediate are shown in Fig. S3.

**Fig. S6.**
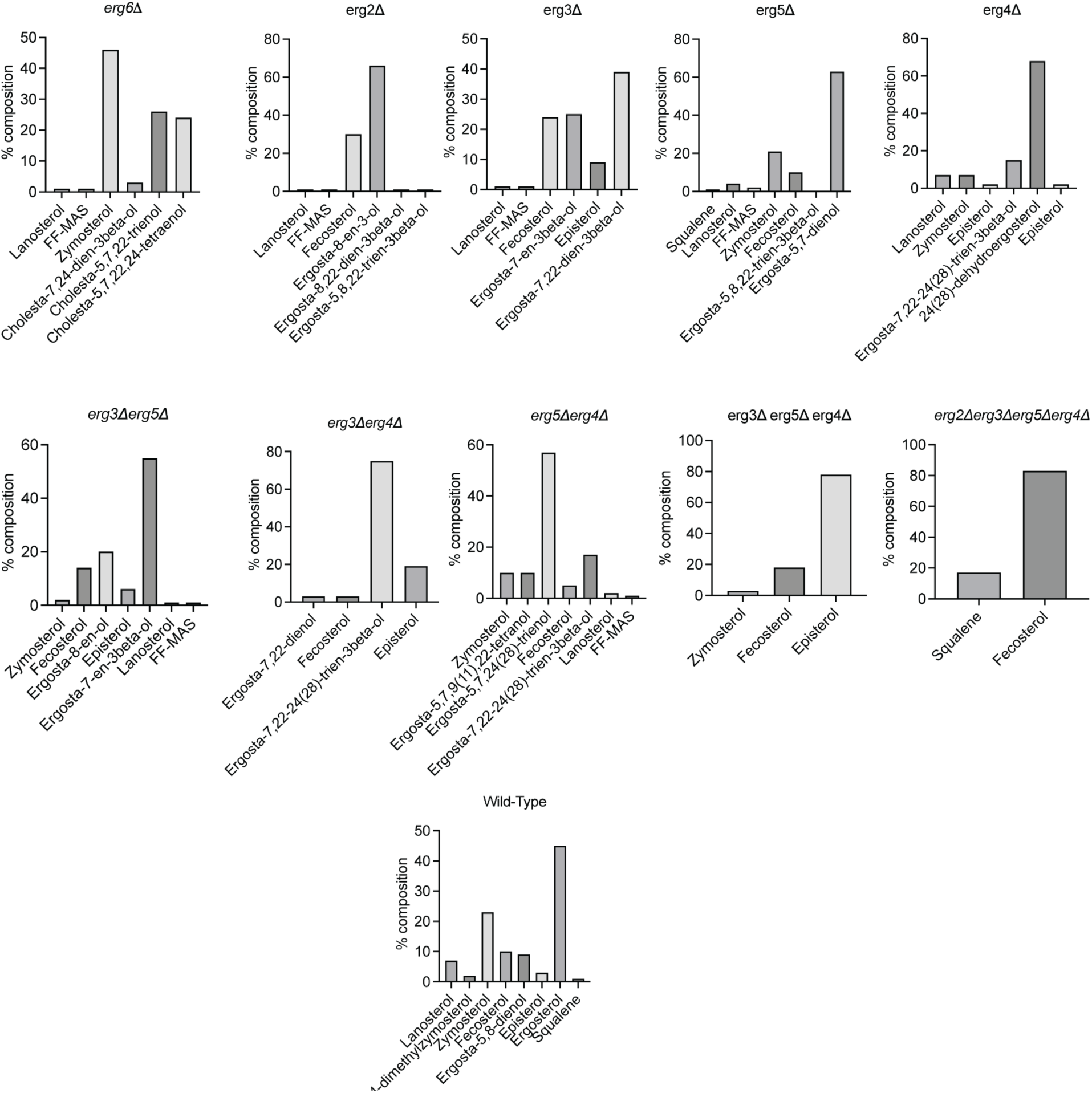
Composition profiles of sterol extracts used for *in vitro* experiments. Abundances were determined by GC-MS.

**Fig. S7.**
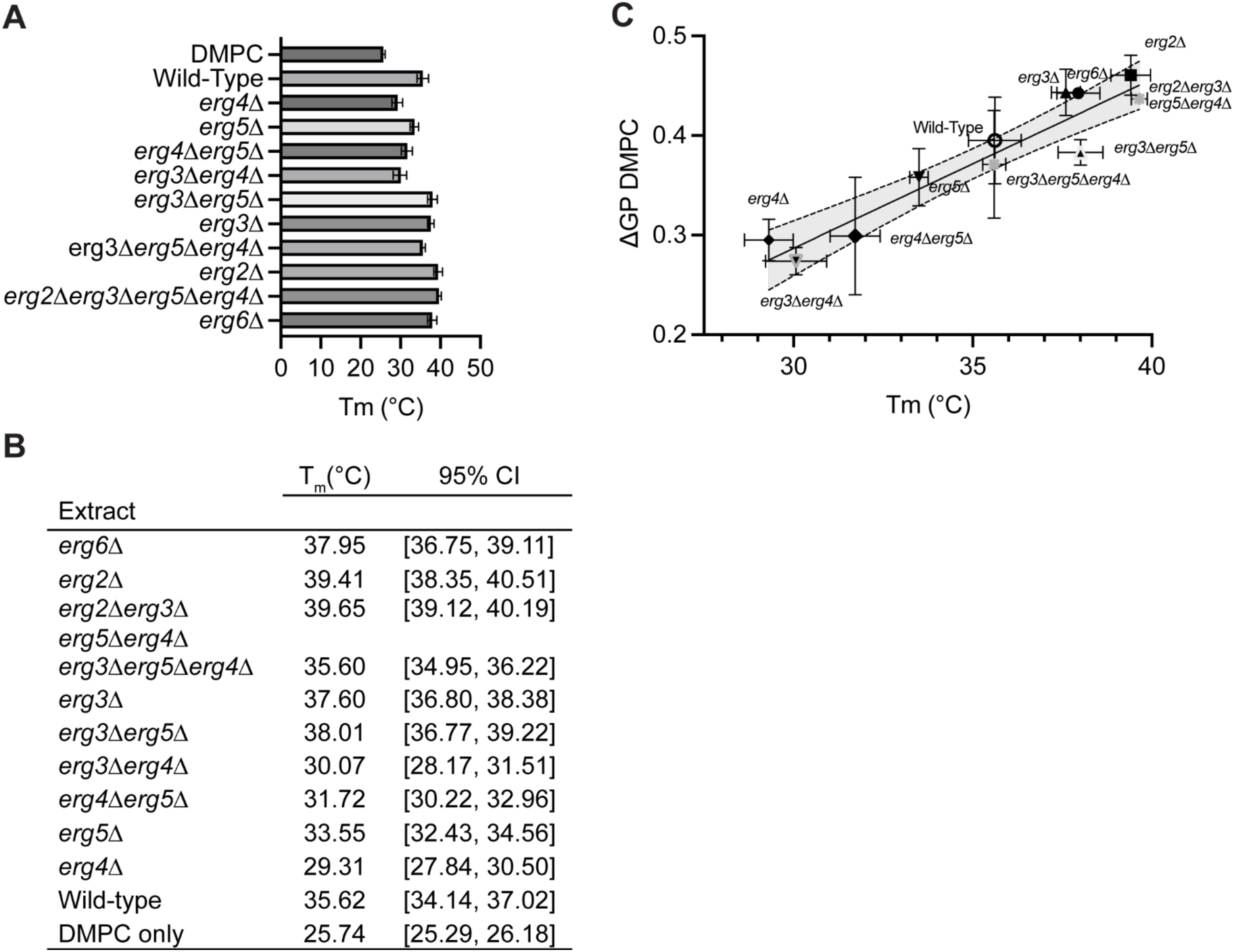
Gene deletion of the initial post-synthetic steps yield sterol extracts with higher transition temperatures compared to deletions of the latter post-synthetic steps. (**A**), Summary of phase transition temperatures of binary mixtures of DMPC and each sterol extract (20 mol %) measured using DPH fluorescence polarization. (**B**), Tabulated summary. (**C**), Regression analysis of the change in Laurdan Generalized Polarization (GP) and phase transition temperature (R^2^ = 0.897, 95% confidence interval shown).

**Fig. S8.**
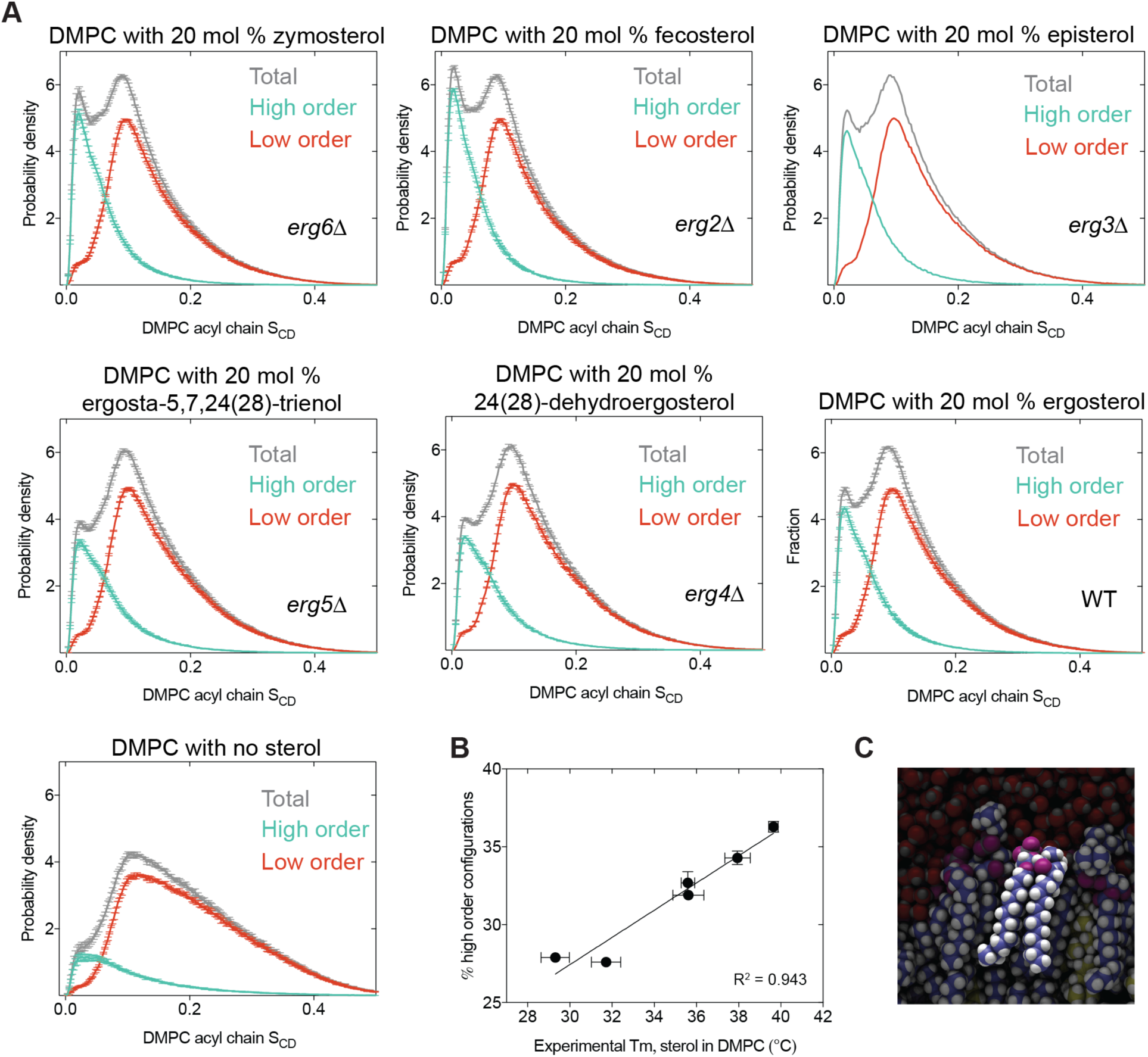
High-order populations in DMPC bilayers. (**A**), Plot of the frequency distribution of all DMPC acyl chain ordering parameters (S_CD_) in simulations for each sterol system, as well as sterol-free DMPC. The distribution shows two distinct peaks: a narrow high order peak on the left, and a broader low-order peak on the right. The cumulative distribution can be deconvolved into these two peaks, with the frequency of the high-order population dependent on the sterol identity in the simulation. Further analysis shows this peak represents DMPC with predominantly *trans* conformers across multiple bonds, while the low-order populations contain mixtures as *trans* and *gauche* conformers. (**B**), Correlation between the proportion of high order configuration in simulations and the experimentally measured Tm for each of the sterol-containing systems. (**C**), Simulation snapshot showing an example of a DMPC molecule with high order configurations of both acyl chains.

**Table S1.**

| Strain | Source | Description |
| --- | --- | --- |
| W303a (*MATa leu2-3112 trp1-1 can1-100 ura3-1 ade2-1 his3-11,15*) | ATCC | Haploid background strain |
| W303a, *erg9::tetO<sub>2</sub>-P<sub>CYC1</sub>-ERG9* | (53) | Repressible sterol metabolism |
| W303a, *erg1::tetO<sub>2</sub>-P<sub>CYC1</sub>-ERG1* | This study | Repressible; accumulates squalene |
| W303a, *erg7::P<sub>MET3</sub>-ERG7* | This study | Repressible; accumulates oxidosqualene |
| W303a, *erg11::tetO<sub>2</sub>-P<sub>CYC1</sub>-ERG11* | This study | Repressible; accumulates lanosterol |
| W303a, *erg24::tetO<sub>2</sub>-P<sub>CYC1</sub>-ERG24* | This study | Repressible; accumulates FF-MAS and derivatives |
| W303a, *erg25::tetO<sub>2</sub>-P<sub>CYC1</sub>-ERG25* | This study | Repressible; accumulates 4,4-dimethylzymosterol |
| W303a, *erg6Δ::his3MX6* | This study | Accumulates zymosterol |
| W303a, *erg2Δ::kanMX4* | This study | Accumulates fecosterol and ergosta-8-en-3-ol |
| W303a, *erg3Δ::his3MX6* | This study | Accumulates predominantly ergosta-5,22-dienol |
| W303a, *erg5Δ::his3MX6* | This study | Accumulates predominantly ergosta-5,7-dienol |
| W303a, *erg4Δ::his3MX6* | This study | Accumulates 24(28)-dehydroergosterol |
| W303a, *erg3Δ::his3MX6, erg5Δ::kanMX4* | This study | Accumulates ergosta-7-en-3-ol |
| W303a, *erg3Δ::his3MX6, erg4Δ::kanMX4* | This study | Accumulates ergosta-7,22,24(28)-trienol |
| W303a, *erg5Δ::kanMX4, erg4Δ::his3MX6* | This study | Accumulates ergosta-5,7,24(28)-trienol |
| W303a, *erg3Δ::leu2, erg5Δ::kanMX4, erg4Δ::his3MX6* | This study | Accumulates episterol |
| W303a, *erg2Δ::trp, erg3Δ::leu2, erg5Δ::kanMX4, erg4Δ::his3MX6* | This study | Accumulates fecosterol |

Yeast (*S. cerevisiae*) strains used in this study.

